# Ocean-scale transcriptomic analysis of corallicolids (Apicomplexa) reveals their ubiquity and molecular interactions with *Pocillopora* corals

**DOI:** 10.64898/2026.05.20.726253

**Authors:** Mathieu Zallio, Clément Leboine, Laura del Rio-Hortega, Maren Ziegler, Alice Moussy, Caroline Belser, Frédérick Gavory, Jean-Marc Aury, Didier Forcioli, Paola Furla, Thamilla Zamoum, Keyla Plichon, Christian R. Voolstra, Clémentine Moulin, Emilie Boissin, Guillaume Bourdin, Guillaume Iwankow, Julie Poulain, Sarah Romac, Tara Pacific Consortium Coordinators, Denis Allemand, Serge Planes, Patrick Wincker, Betina M. Porcel, Quentin Carradec

## Abstract

Corals are complex holobionts, encompassing numerous prokaryotes, viruses, and protists. This associated microbial community strongly influences coral health and its resilience to global ocean warming. Corallicolid apicomplexans are widespread coral–infecting parasites, yet their impact on the coral host remains poorly understood. This knowledge gap largely stems from the low abundance of these parasites in coral tissues, which makes them difficult to isolate and access to their genetic material. Here we analyzed nearly 1,000 *Pocillopora* coral colonies collected from 32 islands during the *Tara* Pacific expedition to identify the drivers of corallicolid prevalence and abundance. Corallicolids were detected in almost all *Pocillopora* colonies with variable relative abundances between islands. The high abundance of specific corallicolid populations correlates with seawater temperature and levels of host protein carbonylation. We used a large collection of 297 metatranscriptomes to assemble a corallicolid transcriptome and we identified apicomplexan parasite signature genes, including the GRA9 and PV2 confirming the close phylogenetic relationship with the family of Eimeriidae. Gene expression patterns indicate that the high abundance of corallicolids correlates with a high transcription of genes encoding apical complex proteins and genes involved in the control of host immune defenses. Overall, this study provides new insights into corallicolid biology and its interaction with the coral host by combining a newly generated transcriptome with a large-scale sampling of *Pocillopora* corals across the Pacific Ocean.

## Introduction

Coral reefs are among the most biodiverse and ecologically important ecosystems on Earth, providing key services such as coastal protection, fisheries, and habitat for approximately one third of marine macroscopic diversity [1]. Yet, these ecosystems are increasingly threatened by global ocean warming, acidification, local pollution, and disease outbreaks [2, 3]. A central challenge for coral reef preservation is to better understand the structure and function of the coral holobiont: the cnidarian host, its main symbiotic microalgae of the Symbiodiniaceae family, the highly diversified bacterial community as well as all other micro- and macro-organisms interacting with the coral colony [4].

Corallicolids are the second most abundant coral micro-eukaryote living within coral tissues [5]. They belong to the apicomplexan phylum, are closely related to coccidians and form a sister lineage to the recently characterized fish parasites ichthyocolids [6, 7]. Interestingly, corallicolids retain nuclear and plastid genes required for chlorophyll-a synthesis despite the absence of functional photosystems [5, 8]. This lineage is therefore an intriguing evolutionary intermediate between photosynthetic free-living chromerids (Vitrella and Chromera) and parasitic apicomplexans such as the human pathogens *Toxoplasma gondii* and *Plasmodium falciparum*.

Corallicolids are mainly present in tropical and subtropical coral reef environments but have also been reported in deep-sea corals and cold waters of various temperate marine ecosystems [9, 10]. Corallicolids are widespread among anthozoans including in the order of Scleractinia, Actinaria, Corallimorpharia, and Octocorallia sub class [5, 9–12]. In several coral populations, the prevalence of corallicolids exceeds 80% [12]. The capacity of a given corallicolid population to infect a narrow or broad range of host species appears to be complex: while specific corallicolid taxa are associated with only one host species, other studies reported the same taxon across diverse hosts or several corallicolid taxa within a single coral colony [13].

Despite the close relationship to many pathogenic taxa within Apicomplexa, corallicolids are frequently found in apparently healthy corals and no clear detrimental effects on the coral host have been observed [5, 12]. Nevertheless, distribution patterns in several corals indicate an opportunistic role. For instance, corallicolids found to be more abundant in diseased than healthy colonies of *Mussismila braziliensis* [14]. More recently, studies on the Mediterranean coral *Paramuricea clavata* and the tropical *Acropora hyacynthus* have shown that corallicolids were relatively more abundant in thermally susceptible than thermally resistant corals [15, 16].

Corallicolid biology, including their potential pathogenicity, remains largely undescribed. This lack of knowledge is, among other, due to the difficulty of accessing corallicolid genetic material. Indeed, corallicolids are not cultivable without their host and their abundance in coral tissue is relatively low, generally below 5% of 18S rRNA sequences of coral associated-microeukaryotes [5, 16]. A recent study developed methodologies to reconstitute corallicolid genes [7]. Specifically, Percoll gradient centrifugations were used to enrich the parasite fraction from two corals: the zoanthid *Parazoanthus swiftii* and the scleractinian *Madracis mirabilis*. Access to multiple nuclear genes was crucial to refine the phylogenetic placement of corallicolids and identify nuclear genes of chlorophyll a biosynthesis. Unfortunately, because the separation between Symbiodiniaceae and corallicolid genes remains challenging, only a minor fraction of the transcriptome could be confidently identified through manual curation of single-gene phylogenies resulting in 185 nuclear genes from Corallicolid ex. *Parazoanthus* and 193 genes from Corallicolid ex. *Madracis*.

Here, we analyzed the corallicolid diversity of nearly one thousand *Pocillopora* colonies sampled across 100 reefs of the Pacific Ocean to study abiotic and biotic factors affecting corallicolid abundance and prevalence. We developed a bioinformatic pipeline to reconstruct a corallicolid transcriptome from 297 *Pocillopora* RNA-seq datasets and get a comprehensive overview of gene expression within the studied system. Using this integrated genetic resource, we characterized corallicolid gene content and get new insights into their biology, functional activity, and interaction with their host.

## Materials and Methods

### *Pocillopora* sampling, sequencing, and identification

Between 2016 and 2018, the schooner *Tara* sampled coral reefs in the Pacific Ocean [17]. At each reef, 10 colonies morphologically identified as *Pocillopora meandrina* were collected, yielding a total of 980 samples [18]. Colonies were sampled at a mean depth of 9.20 ± 3.80 m between 8:30 and 11:30 (local time). Coral fragments were preserved onboard in DNA/RNA shield buffer (Zymo Research) and stored at -20°C prior to nucleic acids extraction as described in Belser et al., 2022. ITS2 and 18SV9 amplicons were sequenced for all *Pocillopora* colonies [20, 21]. In addition, full metagenomes and poly(A)+ transcriptomes were sequenced for 3 colonies per reef. Detailed sequencing protocols are provided in Belser et al., 2022. Species-level identification of *Pocillopora* colonies was performed either using a genome-wide SNP dataset (available for 3 samples per site) or by sequencing a diagnostic marker fragment (available for all samples from I01 to I10 and I15) [22–24].

### Measurements of coral stress biomarkers and seawater parameters

Animal and symbiont biomasses, protein carbonylation levels, and Total Oxygen Scavenger Capacities (TOSC) were measured for five colonies per reef following the methodology described in Porro et al. [25]. Briefly, animal and symbiont biomasses were quantified after NaOH tissue extraction by measuring host protein concentrations using the Bradford assay and Symbiodiniaceae cell densities using hemocytometer counts, with values normalized to coral surface area [26]. Protein carbonylation levels were assessed by ELISA following derivatization of oxidized proteins with dinitrophenylhydrazine, and quantified spectrophotometrically as an indicator of oxidative damage [27]. Finally, Total Oxygen Scavenger Capacities (TOSC) were determined using the TOSC assay, which measures the ability of protein extracts to scavenge peroxyl radicals relative to a Trolox standard [28].

At each reef, physicochemical parameters of the water column were measured *in situ* using multiparameter instruments, including temperature, salinity, dissolved oxygen, pH, nutrient concentrations, and chlorophyll-related variables [18].

### Identification of corallicolid amplicon sequence variants (ASV)

Corallicolid 18SV9 ASVs were identified from metabarcoding ASV tables generated from *Tara* Pacific datasets [20]. ASV tables were filtered to retain ASVs supported by at least 3 reads in one sample. Taxonomic assignment of ASVs was performed using the IDTAXA classifier and similarity-based annotation with VSEARCH, both relying on the PR2 reference database [29–31]. Corallicolid ASVs were defined as those assigned to the Corallicolidae family by at least one of the two methods and were retained for downstream analyses. ASV abundances for each sample were normalised to the total number of eukaryotic 18S rRNA reads. To construct haplotype networks, corallicolid ASVs were aligned using MAFFT v7.490 (G-INS-i algorithm), then inferred using PopART v1.7 and the TCS network algorithm [32, 33].

### Construction of the *Pocillopora* holobiont transcript catalog (PocHTC)

We first removed reads aligned on *Pocillopora cf. effusa* (GCA_942486045.1), *Durusdinium trenchii* (GCA_963969995.1), or *Cladocopium goreaui* (GCA_947184155.2) with >90% identity using BWA-MEM2 v2.2.1 from the 297 RNA-seq datasets [23, 34–36]. Unaligned reads (≥ 100 bp), potentially originating from other organisms of the holobiont, were co-assembled island per island using rnaSPAdes v3.15.5 resulting in a metatranscriptome for each of the 32 islands. These 32 metatranscriptomes were concatenated then clustered using CD-HIT-est v4.8.1 with thresholds of 95 % sequence identity and 80 % coverage [37]. This step removed the redundancy across island-specific assemblies, resulting in a non-redundant transcript catalog for *Pocillopora* holobiont (PocHTC). The coding regions within each transcript were predicted with TransDecoder v5.5.0 [38].

### PocHTC taxonomic affiliation and abundance levels

All transcripts were aligned against a concatenated protein database combining NCBI nr version 08-2023 and MarFERReT version 1.1 databases using DIAMOND blastx v2.1.2 with the *--sensitive* option [39, 40]. For each transcript, the best hit with ≥40% identity and ≥50 amino acids was retained. The complete taxonomic lineage of nr hits was retrieved from the NCBI taxdump (version 08-2023). To remove remaining *Pocillopora* transcripts from the catalog, we aligned PocHTC transcripts against available *Pocillopora* reference genomes (GCA_014529365.1, GCA_036669915.1, GCA_942486045.1, GCF_030620025.1, and GCF_003704095.1) using NCBI BLASTn v2.16 [41]. Transcripts aligned with ≥ 90% sequence identity over ≥150 bp alignment length were excluded. RNA-seq and genomic reads of the 297 *Pocillopora* colonies were mapped to the PocHTC using BWA-MEM2 v2.2.1. Reads ≥100 bp aligned with ≥95% identity over ≥50% of their length were selected to estimate abundance and expression levels. Raw counts were obtained using samtools idxstats then normalised to transcript per million (TPM).

### Elucidation of a corallicolid reference transcriptome

Transcripts detected in at least five samples were sorted into transcriptome bins using CONCOCT v1.1.0 [42], which combines abundance profiles across samples and tetranucleotide frequencies to group transcripts likely originating from the same organism. A minimum transcript length of 300 bp was applied. The taxonomy of each transcriptome bin was determined based on previously computed taxonomic affiliation of the transcripts. Transcriptome bins containing more than 7,000 transcripts and showing ≥ 30% completeness estimated with BUSCO v5.2.2 [43] were selected. The appropriate BUSCO lineage dataset was selected according to the bin taxonomy. For the corallicolid transcriptome bins, completeness was assessed using the *alveolata_odb10* database. Corallicolid transcripts were translated in full on the 6 frames using EMBOSS Transeq v6.6.0 to search for protein domains using InterProScan v5.61.93.0, DIAMOND Blastx on the KEGG protein database and KofamScan v1.3.0 [39, 44, 45]. Samples with less than 2,000 expressed corallicolid genes were excluded from transcriptomic analyses. Gene expression levels of corallicolid transcripts were normalised in TPM to be comparable between samples whatever the sequencing depth or the corallicolid abundance in the metatranscriptome.

### Corallicolid phylogenetic analyses

Published Apicomplexa 18S rRNA sequences ≥800 bp containing the V9 region were retrieved from Trznadel et al., 2024. The 18S rRNA sequence from an Ichthyocollid was included as the closest outgroup to Corallicolida [6]. Sequences were aligned using MAFFT v7.490 with the G-INS-i algorithm [46]. The alignment was curated with goalign v0.3.1 by removing positions with ≥50% gaps. An initial maximum-likelihood tree was inferred with IQ-TREE v3.0.1 to identify the best-fitting substitution model using ModelFinder Plus. A final reference tree was reconstructed with RAxML-NG v1.1.0 under the TIM3+F+G4 model, with branch support estimated from 1,000 bootstrap replicates [47]. Environmental ASVs were aligned onto the reference alignment using PaPaRa v2.5 [48]. Phylogenetic placement was performed with EPA-NG v0.3.8 [49]. Placement results were analyzed using GAPPA v0.9 to compute likelihood weight ratios (LWR) and expected distances between placement locations (EDPL) [50]. Tree visualization and metadata integration were conducted using Anvi’o v8 [51].

For the phylogeny based on nuclear genes, representative apicomplexan species from Coccidians, Protococcidians, Ichthyocholids and *Plasmodium falciparum* were included. A NCBI BLASTp search was performed between 115 previously identified conserved proteins [7] and the Corallicolid ex. *Pocillopora* proteome. The best hit per gene with an e-value < 1e-5 was retained. Multiple alignments were computed with MAFFT L-INS-i version 7.490 then concatenated [46]. Concatenated multiple-alignments were trimmed with trimAl –gt 0.6 version 1.5.0 [52], then a phylogeny was inferred with RaxML using the PROTGAMMALG model and 100 rapid bootstrap replicates [47]. Branch support was calculated with the TBE normalisation method implemented in BOOSTER [53], and trees were visualised in iTOL [54].

### Orthology analyses

An orthology analysis was performed with OrthoFinder (v3.0) [55] on the predicted proteome of Corallicolid ex. *Pocillopora*, 66 parasitic apicomplexans, and two chrompodellids retrieved from the VEuPathDB (Release 68) database [56]. Protein domains and functional signatures of selected OGs were annotated using InterProScan (v5.75.106.0) [57]. Conserved motifs were identified with MEME (v5.5.7) in protein mode, allowing up to 10 motifs with an e-value < 1e-5. For phylogenetic analysis, protein sequences were aligned with MAFFT (v7.490) using the FF-NS-i algorithm [46, 58]. The alignment was then filtered to remove sites containing more than 50% gaps using Goalign (v0.3.1) [59]. Phylogenetic reconstruction was performed with IQ-TREE (v1.6.12), enabling automatic model selection using ModelFinder Plus [60]. Node support was assessed with 1000 ultrafast bootstrap replicates and 1000 SH-aLRT replicates. The final tree was visualised and annotated using iTOL (v7.2.2) [54].

For the structural analysis of GRA9 protein, six representative reference proteins were selected: 3 Sarcocystidae (*Besnoitia besnoiti*, *Cystoisospora suis*, *Toxoplasma gondii*), 2 Eimeriidae (*Eimeria acervuline* and *Eimeria tenella*) and the predicted corallicolid homolog. For the reference species, 3D protein structures were retrieved directly from the AlphaFold Protein Structure Database, using the available models without modification. Corallicolid protein structures were predicted using the AlphaFold Protein Structure Prediction online server (https://alphafoldserver.com/) [61]. All structures were subsequently superimposed in PyMol (v3.1.6.1) using the CEalign algorithm for optimal local-geometry alignment [62]. The structural distance (RMSD) was computed in PyMOL. Global structural similarities between pairs of proteins were quantified using TM-score, calculated pairwise with the online TM-align tool (https://aideepmed.com/TM-align/) following default parameters [63].

## Results

### Diversity and prevalence of corallicolids in *Pocillopora* corals

During the *Tara* Pacific expedition, 980 *Pocillopora* colonies belonging to 6 species were collected across the Pacific Ocean (Fig. 1A). The V9 region of 18S rRNA sequences was sequenced, generating an amplicon sequence variant (ASV) table encompassing all eukaryotic ASVs of the holobiont [20]. Within the Apicomplexa phylum, 34 ASVs were assigned to corallicolids based on available Corallicolid sequences of the PR2 database [29]. These ASVs were further analyzed to investigate the diversity, abundance and prevalence of corallicolids in *Pocillopora* colonies. Corallicolid ASVs formed a monophyletic group with previously identified corallicolid 18S rRNA sequences and were closely related to the recently described Ichtyocolid lineage (Fig. S1). These ASVs were highly similar differing by only a few base pairs (Fig. 1B). Among the corallicolid ASVs, 6 showed a high prevalence (above 20%) across all *Pocillopora* colonies, with ASV-3 reaching a prevalence of 92.3%. These 6 ASVs also displayed the highest mean relative abundance, with ASV-1 reaching up to 0.34% of all eukaryotic ASVs (Fig. 1B and Fig. S2A). Corallicolid were not detected in only 7 *Pocillopora* colonies collected in Upolu (I10), Kiribati (I13) and in the Gulf of California (I30) despite similar sequencing depth. Overall, these results indicate that corallicolids are nearly ubiquitous in *Pocillopora* corals throughout the Pacific Ocean.

**Fig. 1.**
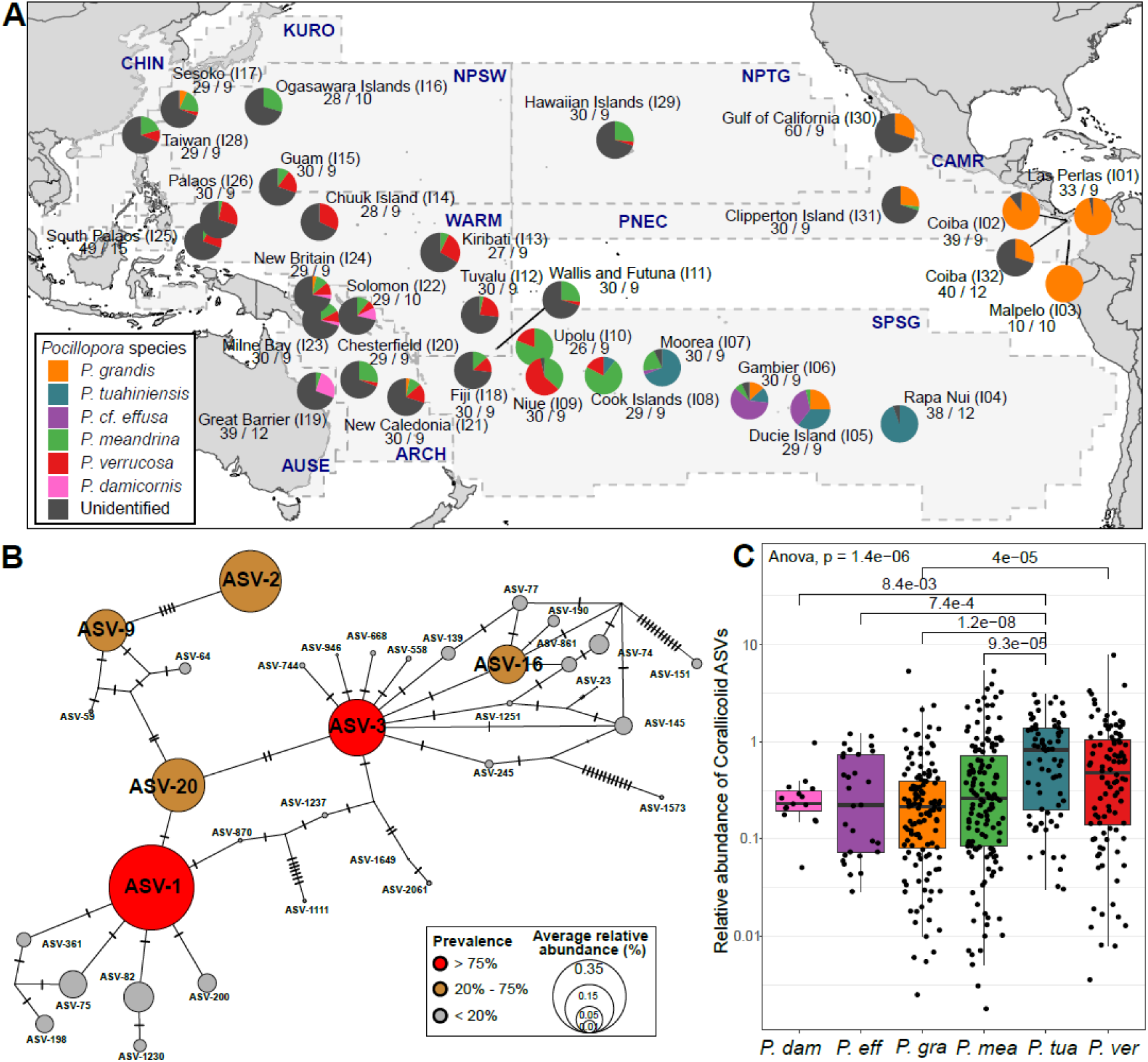
*Pocillopora* sampling map and corallicolid diversity. A) *Pocillopora* sampling during *Tara* Pacific expedition. Circular diagrams represent different species collected around each island. *Pocillopora* colonies for which the species has not been identified are in grey. The number of individuals with available 18SV9 rRNA amplicons (left number) and available metatranscriptomic datasets (right number) are indicated under each diagram. Longhurst provinces are delimitated by dashed lines and their name codes are in blue. B) TCS haplotype network of corallicolid 18SV9 rRNA sequences. Corallicolid ASV sequences are represented by circles displayed in the network according to their number of nucleotide variants. Each nucleotide difference between 2 ASV sequences are represented by small dashes. Circle size is proportional to the average relative abundance of the ASV among all 18SV9 sequences across all *Pocillopora* colonies. Prevalence across *Pocillopora* colonies is indicated by circle colour. C) Distribution of corallicolid ASV relative abundance across the 6 *Pocillopora* lineages. Significant statistical differences in mean abundances (Wilcoxon p.values <0.01) are indicated for each pair of species.

ASV-1 and ASV-3 were the dominant corallicolid sequences in 85% of *Pocillopora* colonies (Fig. S2A). Interestingly, the ASV-1 sequence was 100% identical to a corallicolid previously identified in the actiniarian sea anemone *Metridium senile* (OR883653), while ASV-3 was 100% identical to corallicolids identified in the corallimorpharians *Corynactis* spp. (OR883651, OR883652) and *Rhodactis* sp. (MH304760; Fig. S1). These studies sampled corals from cold waters of British Columbia, suggesting that these corallicolids occupy a wide range of hosts and environmental conditions.

Although the mean relative abundance of corallicolids was low, it ranged from 0 to 9.25% of total ASV reads in individual colonies (Fig. 1C). We observed a significantly higher average abundance in *Pocillopora tuahiniensis* (0.91%, SD = 0.78) compared to *Pocillopora damicornis* (0.29%, SD = 0.21), *Pocillopora cf. effusa* (0.40%, SD = 0.37), *Pocillopora grandis* (0.36% SD = 0.59), and *Pocillopora meandrina* (0.59%, SD = 0.86). ASV-1 was significantly more abundant in *P. tuahiniensis, P. grandis*, *P. verrucosa*, and *P. meandrina*, whereas ASV-3 was more abundant in *P. cf. effusa* (Fig. S2B). The 4 other abundant ASVs were dominant only in a limited number of colonies of *P. verrucosa*, *P. meandrina*, and *P. cf. effusa* (Fig. S2B). ASV-1 and ASV-3 co-occurred in 256 *Pocillopora* colonies (26% of the dataset), and the 4 most abundant ASVs (ASV-1,2,3,20) co-occurred in 17 colonies (1.7% of the dataset; Fig. S3).

*Cladocopium* and *Durusdinium* are two Symbiodiniaceae genera commonly associated with *Pocillopora* corals. In our dataset, 61 out of 126 *P. grandis* and 21 out of 76 *P. verrucosa* colonies were associated with *Durusdinium* [64]. No significant differences in corallicolid ASV diversity or abundance were detected according to the Symbiodiniaceae genus (Wilcoxon test p-values=0.57 for *P. grandis* and 0.74 for *P. verrucosa*; Fig. S2C and S2D). Additionally, no correlation was observed between the relative abundances of corallicolids and Symbiodiniaceae (Fig. S4). The number of ASV reads for Symbiodiniaceae were generally far more abundant than for the corallicolids, with only 7 exceptions in which both symbionts had similar ASV relative abundances in *Pocillopora* colonies (Fig. S4).

### Drivers of corallicolid abundance in the Pacific Ocean

The relative abundance of corallicolid ASVs strongly varied among sampled islands (Fig. 2A). *Pocillopora* colonies at the Hawaiian Islands (I29) showed the highest corallicolid abundance across the entire Pacific Ocean, with a median of 2.28% of total ASV reads. Moorea (I07) and Niue (I09) islands exhibited the highest corallicolid abundances of the South Pacific province (SPSG), dominated by ASV-1. In the Coral Triangle (I22-I26), corallicolids are also abundant with the higher ASV diversity. Interestingly, 6 islands (I03, I05, I10, I12, I13, and I30) display consistently low corallicolid relative abundances (<0.1%; Fig. 2A). Redundancy analyses (RDA) were performed using *in situ* environmental parameters to investigate the drivers of corallicolid diversity and abundance (Fig. 2B and S5). ASV-3 abundance was correlated with low seawater temperature of the Northwest Pacific (NPSW) and Rapa Nui (I04) islands (Spearman = 0.36, p-value < 1e-5), whereas ASV-1 was associated with warmer environments (Spearman = -0.44; p-value < 1e-5; Fig. S5).

**Fig. 2:**
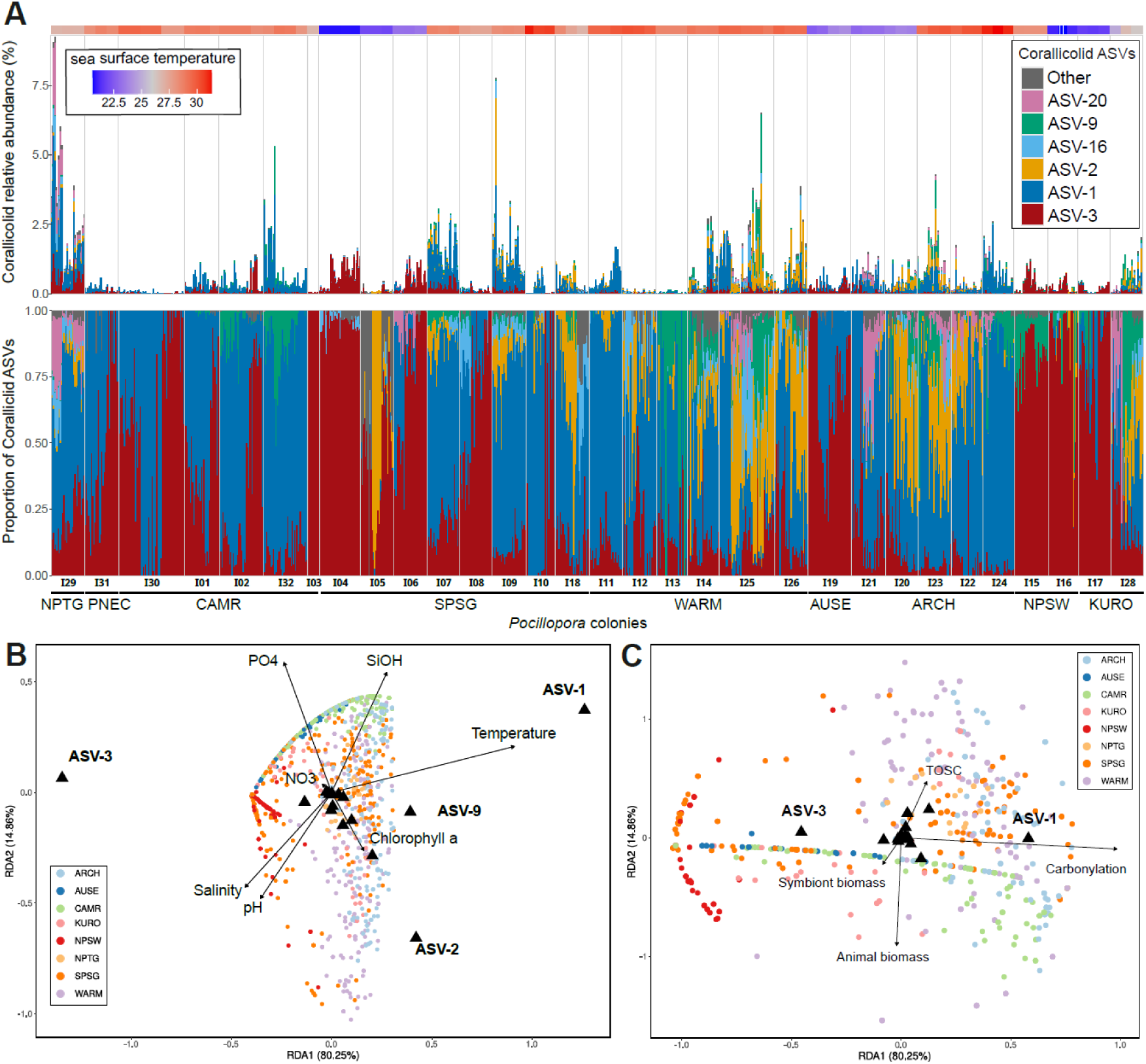
Drivers of corallicolid ASV diversity and abundances. **A)** ASV abundance relative to the total number of 18SV9 sequences from each coral colony (top panel) or relative to the total corallicolid abundance (bottom). The 6 most abundant corallicolid ASVs are represented. The average sea surface temperature of each sampled reef is indicated above each bar. **B and C)** RDA of corallicolid ASVs diversity according to 6 environmental parameters (Temperature, SiOH, PO4, Salinity, pH and Chla) (B) or 4 coral host stress markers: animal biomass, symbiont density, carbonylation and TOSC (antioxidant defenses; C). *Pocillopora* samples are represented by dots coloured by oceanic provinces and the main corallicolid ASVs by black triangles.

To understand whether the presence of corallicolids correlates with host physiology, we used 4 stress biomarkers measured on the same colonies [25]. Animal and symbiont biomass are representative of polyp tissue thickness and symbiont density respectively. The redox state of *Pocillopora* colonies is quantified by the rate of carbonylated proteins and the antioxidant defenses by the Total Oxidant Scavenging Capacity (TOSC; see methods). ASV-1 abundance is weakly but significantly correlated with levels of protein carbonylation (Spearman = 0.21; p-value = 6.4e-6), whereas ASV-3 negatively correlated with protein carbonylation (Spearman = -0.25; p-value = 1.8e-7; Fig. 2C, S5).

### Reconstruction of a corallicolid transcriptome and gene expression patterns

We developed a bioinformatic pipeline to reconstruct corallicolid genes and investigate their activity across the sampled islands. We used 297 metatranscriptomes from *Pocillopora* colonies to assemble all genes of the holobiont after excluding coral host and Symbiodiniaceae reads based on sequence homologies. We obtained a transcript catalog of the *Pocillopora* holobiont (PocHTC) containing 2.8 million transcripts. To identify corallicolid transcripts within this catalog, we clustered PocHTC transcripts using their abundances and their tetranucleotide frequencies (see Methods). Among the reconstructed transcriptomes, we obtained a cluster taxonomically affiliated to Conoidasida (Apicomplexa) containing 13,802 transcripts, and being 33.4% complete according to the BUSCO score.

We determined the phylogenetic position of this transcriptome, by using recently sequenced corallicolid nuclear genes from the golden zoanthid, *Parazoanthus swiftii*, and a yellow pencil coral *Madracis mirabilis* [7]. We obtained significant matches for 148 out of 193 Corallicolid ex. *Madracis* proteins and 141 out of 185 Corallicolid ex. *Parazoanthus* proteins. Corallicolid ex. *Pocillopora* had more than 12% divergence with previously sequenced corallicolids. We performed a phylogenomic analysis based on the concatenation of 102 proteins, previously selected to reconstruct an Apicomplexan tree [7]. The phylogeny confirms the phylogenetic position of corallicolids as being closely related to ichtyocolids and reveals that, despite substantial divergence, the 2 corallicolids identified in scleractinian corals are more closely related to each other than to the corallicolid identified in the zoanthid (Fig. S6). Interestingly, none of the 3 nuclear genes involved in chlorophyll a biosynthesis identified in Corallicolid ex. *Parazoanthus* (ChlI, ChlM and ChlG) were detected in the Corallicolid ex. *Pocillopora* transcriptome. Although this absence could be due to the incomplete transcriptome, we cannot exclude the possibility that these genes were not retained in the nuclear genome of this corallicolid.

To link the transcriptome to the corallicolid ASVs, we performed Spearman correlations between the abundances of each of the 34 corallicolid ASVs and the transcriptome abundance. The highest correlation was observed for ASV-3 (Spearman = 0.81: p-value < 0.01), followed by ASV-1 with a lower score (Spearman 0.47, p-value < 0.01; Fig. 3A). We selected the 34 *Pocillopora* samples in which at least 2,000 corallicolid transcripts were detected, we computed the relative expression levels of each of the 13,802 corallicolid transcripts in each sample, and used these normalized counts to identify drivers of gene expression patterns across 8 Pacific islands. We observed a strong biogeographic signal, with samples collected from the same island displaying similar expression profiles (Fig. 3B). Interestingly, despite the relatively close geographic proximity the 3 sites sampled around Hawaii (I29) exhibited distinct corallicolid expression profiles. Corallicolid genes in different *Pocillopora* species from the same island presented very similar expression profiles (e.g. *P. damicornis* and *P. meandrina* in I19; *P. grandis* and *P. cf. effusa* in I06). This result suggests that the host species within the same genus has a limited impact on corallicolid activity. Finally, very distant islands such as the Great Barrier Reef (I19), Las Perlas (I01), and Ogasawara islands (I16) exhibited similar corallicolid gene expression profiles, indicating that geographic distance alone is not sufficient to explain the observed expression patterns (Fig. 3B). In addition, distinct corallicolid gene expression patterns were associated with a low level of protein carbonylation in Rapa Nui Island (I04) and a higher activity of antioxidative defenses (TOSC) in the Hawaiian Islands (I29; Fig. 3C). These islands are among those with a high relative abundance of corallicolids (Fig. 3A).

**Fig. 3:**
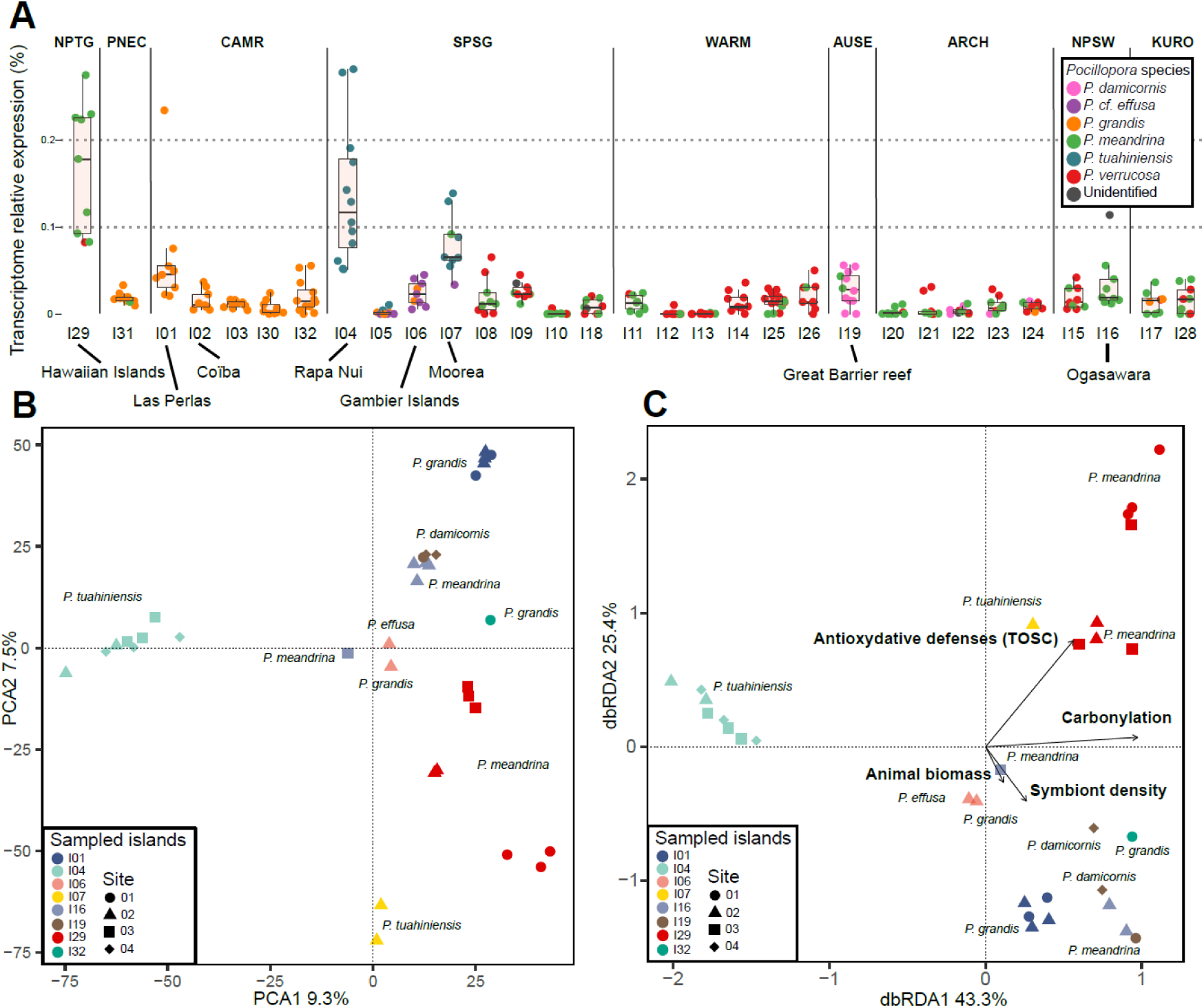
Gene expression patterns of the corallicolid transcriptome across *Pocillopora* samples. A) Relative corallicolid transcriptome expression in each island. Each dot is the ratio of the sum of the expression of all genes relative to all sequenced metagenomic reads of each sample. The boxplots show the median, 1^st^ and 3^rd^ quartiles of the distribution per island. The 6 *Pocillopora* species are represented by dot coulours. B) Principal Component Analysis of gene expression levels in 34 *Pocillopora* colonies, in which more than 2,000 corallicolid genes are expressed. The different shapes indicate the different reefs sampled on each island and the *Pocillopora* species are indicated. C) Distance-based redundancy analysis (db-RDA) using stress biomarkers, host biomass and symbiont density as variables.

### Expressed gene functions in different coral reefs

We predicted putative functions on 3,478 transcripts based on InterProScan and KEGG analyses. Among the most highly expressed transcripts with assigned functions, we identified 7 transcripts involved in glycolysis, including fructose-bisphosphate aldolases, glyceraldehyde-3-phosphate dehydrogenases, enolases, and phosphoglycerate kinases (Fig. 4A). The pronounced expression of genes involved in this pathway indicates corallicolids to be metabolically active and suggests that carbohydrate catabolism represents a major energy source. We also detected 179 transcripts encoding ribosomal proteins, reflecting high translational activity.

**Fig. 4:**
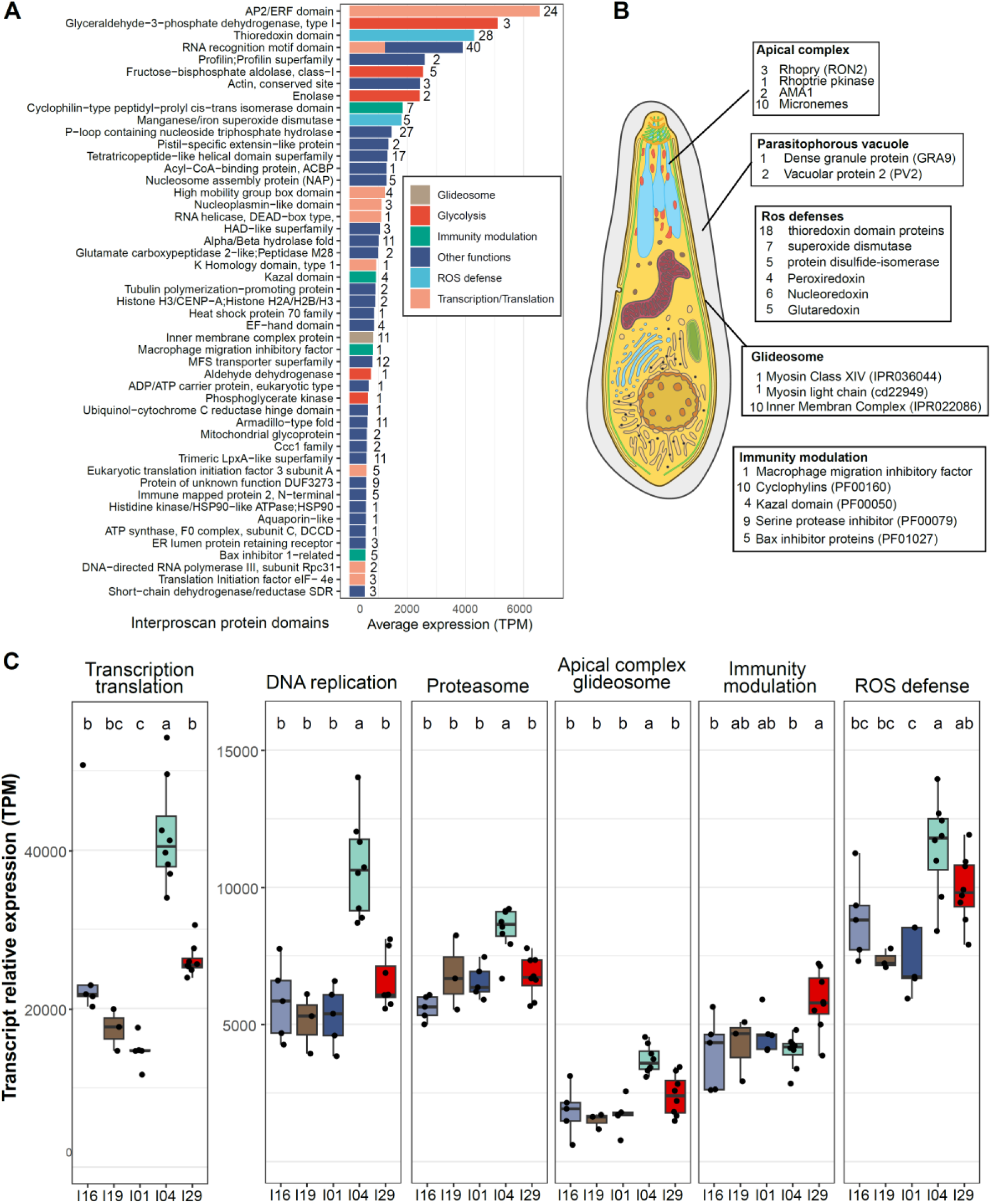
Expressed gene functions in corallicolids. A) The 50 most expressed protein functional domains are shown and coloured according to their functional categories. The number of transcripts carrying each domain is indicated on the right of each bar. Gene expression was normalised in TPM across the 34 samples in which more than 2,000 corallicolid genes are detected. Transcripts encoding ribosomal proteins and those of unknown functions were excluded from this panel. B) Schematic representation of an apicomplexan [76]. Key proteins involved in parasite infection and detected in Corallicolid ex. *Pocillopora* are indicated. C) Relative expression of corallicolid transcripts involved in different biological processes. Islands with at least 3 samples expressing more than 2,000 expressed genes are represented. Significant differences in mean expression levels based on Tuckey’s post hoc tests are indicated above each boxplot in letter codes.

Several transcription factors known to play key roles in *P. falciparum* and *T. gondii* activity were identified, including proteins containing the AP2/ERF domain, the high-mobility group box domain, the nucleoplasmin-like histone chaperone domain, and the K homology domain type 1 [65–68]. A total of 11 transcripts encoding inner membrane complex (IMC) proteins were detected in Corallicolid ex. *Pocillopora* (Fig. 4A and B). These proteins, localized to a double-membrane under the plasma membrane in apicomplexan parasites, are thought to be important for motility and host invasion [69].

Among the most highly expressed transcripts, we identified a Glutathione peroxidase, a thioredoxin, and an iron superoxide dismutase, suggesting a strong antioxidant activity and an active response to limit oxidative stress (Fig. 4A, 4B). Proteins potentially involved in modulating host immunity were also highly expressed, including the macrophage migration inhibitory factor (MIF), a phenylpyruvate tautomerase known to modulate host cytokine activity in *P. falciparum* [70, 71]. In addition, we identified 10 putative cyclophilins, 4 transcripts carrying the Kazal domain, which are known to protect against host digestive enzymes in *T. gondii*, and 9 serine protease inhibitors (SerPIN) reported in several parasites to limit the host immune responses [72–74]. Finally, 5 Bax inhibitor proteins were highly expressed. Downregulation by *Eimeria* and *T. gondii* of host pro-apoptotic Bax proteins has been shown to be important to ensure parasite proliferation before host cell death [75].

We then compared the expression levels of genes involved in major biological processes of Apicomplexa across several Pacific islands (Fig. 4C). Genes involved in transcription, translation, DNA replication, proteasome activity, as well as genes of the apical complex and glideosome were significantly higher expressed in Rapa Nui (I04) than in all other sampled islands. This result indicates a higher activity of corallicolids in this region, potentially explaining the high relative abundance of ASV-3 (Fig. 2A). Interestingly, genes involved in host immune modulation were less expressed in Rapa Nui (I04) than in Hawaiian Islands (I29), suggesting that corallicolids around Rapa Nui (I04) did not elicit a strong response to the host immune system. In both islands, genes involved in reactive oxygen species (ROS) defence were higher expressed than in islands with lower corallicolid abundance (Fig. 4C). Overall, these results reveal distinct gene expression strategies that may influence the efficiency of apicomplexan invasion and persistence in coral tissues.

In addition to the most highly expressed functions, several genes potentially involved in host-parasite interaction were examined in greater detail. We therefore performed a homology-based approach on our predicted corallicolid proteins and those of 66 reference apicomplexan parasites covering several clades (Gregarines, *Cryptosporidium*, Haemosporida, Piroplasmida, and Eimeriida). Two non-parasitic chrompodellidae species (*Chromera velia* and *Vitrella brassicaformis*) were also included. A total of 5,407 corallicolid proteins were clustered into 2,182 orthogroups (OGs), of which 586 contained only parasitic sequences and no chrompodellid proteins. Among these, 395 OGs were specific to corallicolids, whereas 191 were shared with reference apicomplexans (Fig. S7). While some of these 191 shared OGs included proteins from many apicomplexan lineages, others were restricted to only a few clades, reflecting both a conserved genetic core among apicomplexans and lineage-specific evolutionary trajectories. These shared OGs include proteins involved in host-parasite interactions and invasion, such as micronemal (MIC), rhoptry neck (RON), parasitophorous vacuole (PV) and dense granule (GRA) proteins, as well as proteins associated with cytoskeletal organisation and parasite motility, including IMC proteins.

Nearly half of the 191 OGs shared with other apicomplexans (84 OGs) were restricted to the order Eimeriida (Eimeriidae and Sarcocystidae; Fig. S7). These OGs were enriched in proteins previously reported to play key roles in parasitic interactions with the host, such as invasion and motility (e.g., GRA9 and PV2), highlighting a shared evolutionary history of these proteins within this clade.

### Evolution of GRA9 and PV2 proteins in Eimeriida

GRA9 is a dense granule secreted protein released into the parasitophorous vacuole following host cell invasion and is known to play a key role in maintaining the host-parasite interface and facilitating nutrient acquisition in *Toxoplasma gondii*. To assess whether corallicolid encode a GRA9 homolog, we analyzed the OG (OG0003106) containing the putative GRA9 protein from corallicolid as well as those from other apicomplexans. This OG was composed of two different proteins for most reference species: one unannotated protein (generally > 1000 aa) and one protein of approximately 300 aa, a size corresponding to GRA9 already described for apicomplexan species [77]. Protein motif analysis revealed distinct patterns, conserved at the family level (Fig. 5A). The corallicolid protein exhibited a distinctive pattern composed of motifs present in both Eimeriidae and Sarcocystidae families in the same order (Fig. 5A). This result was consistent with the phylogenetic GRA9 reconstruction, which placed the corallicolid protein basally to all other GRA9 sequences. Structurally, the corallicolid GRA9 protein belongs to the same structural family of other apicomplexan GRA9 (TM-score 0.5-0.6, RMSD 3.2-3.6 Å) and is equally distant from Sarcocystidae and Eimeriidae, with peripheral differences corresponding to regions of lower structural confidence (pLDDT < 50) (Fig. S8). Overall, the pattern composition, phylogenetic analysis, and structural similarity strongly suggest that the protein belonging to corallicolid is indeed a homolog of GRA9. These findings highlight the presence of a conserved host-parasite interaction protein conserved in corallicolids, maintained across Eimeriida despite extensive evolutionary divergence.

**Fig. 5:**
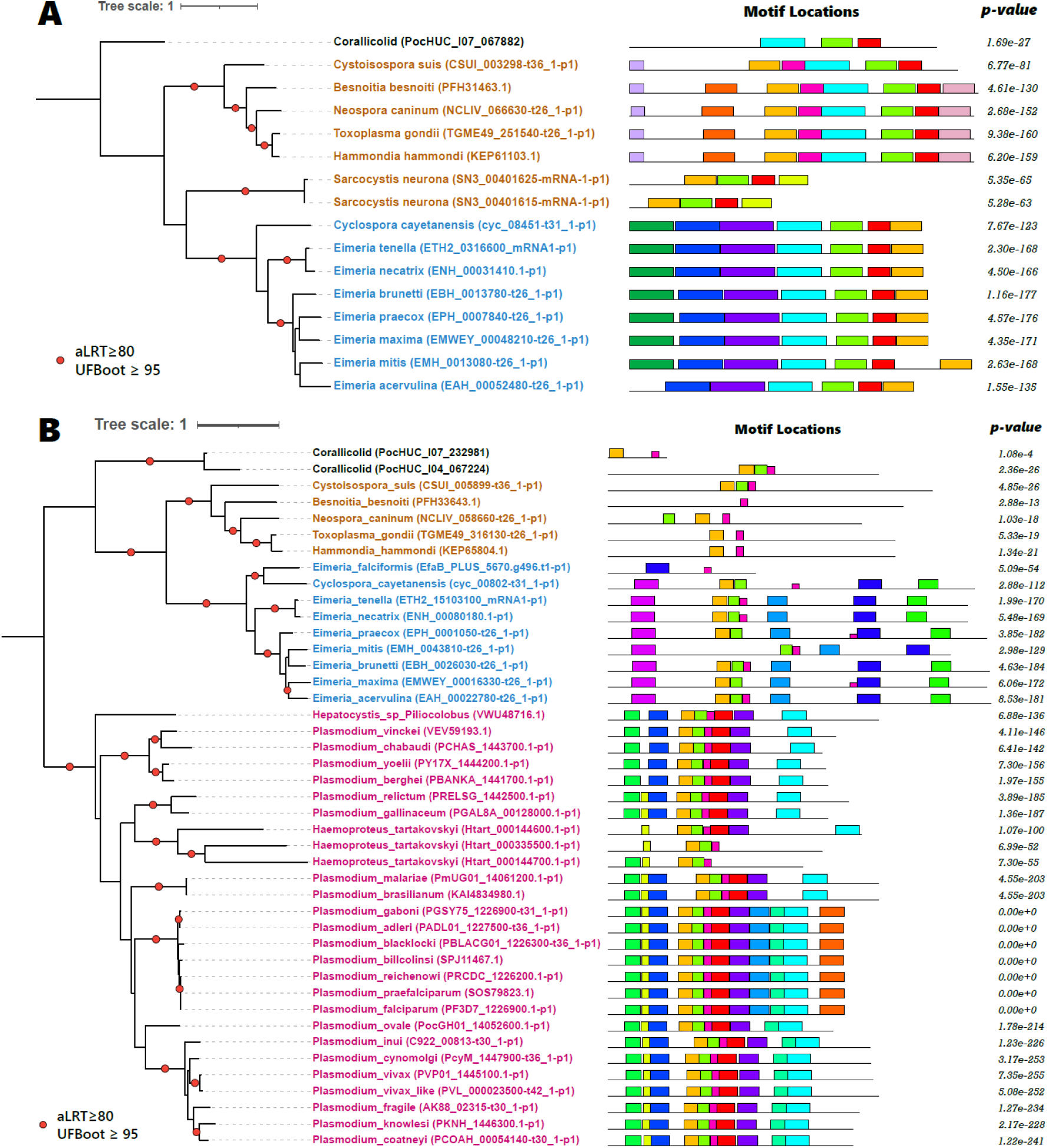
Phylogenetic relationships and motif conservation of apicomplexan GRA9 and PV2 proteins. (A) Maximum-likelihood phylogeny of GRA9 proteins extracted from the corresponding OG. (B) Maximum-likelihood phylogeny of PV2 proteins from the corresponding orthogroup. Phylogenetic trees were inferred with WAG+F+I+G4 model for GRA9 and JTT+F+R3 model for PV2. Nodes with SH-aLRT ≥ 80 and UFBoot ≥ 95 are indicated. Branches are colored according to apicomplexan clades, with corallicolid sequences shown in black, Eimeriida in blue, Sarcocystidae in orange, and Haemosporida in pink. For each protein, conserved sequence motifs are shown alongside the trees with colored boxes Associated motif significance values (p-values) are reported on the right.

We also investigated the presence of other parasitophorous-vacuole associated proteins in corallicolids, focusing on the OG containing proteins annotated as parasitophorous vacuolar protein 2 (PV2) in Haemosporida (OG0002569). This OG was exclusively composed of sequences from Eimeriida, Sarcocystidae and Haemosporida, with no representatives outside these clades. The corallicolid encoded two proteins of 532 and 125 aa within this OG, the longest exhibiting a size comparable to PV2 proteins from Haemosporida and Sarcocystidae. Motif analyses revealed lineage-specific architecture patterns and 3 conserved motifs across all parasitic clades including the corallicolid (Fig. 5B). Finally, the phylogeny based on PV2 proteins shows corallicolid PV2 homolog branching basally relative to the Haemosporida and Sarcocystidae clades, and well-supported family-level groupings (aLRT ≥ 80 and UFBoot ≥ 95). Together, these results indicate that corallicolids encode a divergent PV2 protein, retaining a minimal conserved core shared across parasitic Apicomplexa. This finding supports the hypothesis that key components of the parasitophorous vacuole were already present in the ancestor of parasitic intracellular Apicomplexa and have subsequently undergone lineage-specific remodeling.

## Discussion

The large number of *Pocillopora* samples analyzed in this study allowed us to identify robust patterns of corallicolid prevalence, abundance and activity across the Pacific Ocean. Corallicolids were found to be nearly ubiquitous in *Pocillopora* corals and exhibited low genetic diversity, with only 2 dominant ASVs, 4 minor ASVs, and 30 rare variants. Although, corallicolid relative abundances were generally low (mean 0.34%), they varied strongly among reefs, which emerged as the primary driver of abundance patterns. Despite their relatively low abundance in *Pocillopora*, we successfully reconstructed a partial corallicolid transcriptome using *de novo* metatranscriptome assembly followed by co-abundance clustering. Gene expression analyses revealed distinct transcriptomic profiles across sampled islands. Elevated expression of genes involved in host immune modulation in the Hawaiian Islands and increased activity of the apical complex and glideosome in Rapa Nui correlated with higher corallicolid abundance in these regions. Together, these results provide the first insights into the biological interactions between corallicolids and their coral hosts.

### Construction of a corallicolid reference transcriptome based on metagenome co-abundances

Since the discovery of an unknown apicomplexan lineage commonly associated with corals [78], the nature of corallicolid-coral interactions remains unresolved. While sequencing of the corallicolid plastid genome in 2019 clarified the phylogenetic position of this lineage [5], access to its full genome or transcriptome content has remained a critical gap.

Recovering corallicolid genetic material is particularly challenging because these parasites occur within complex coral holobionts comprising several eukaryotes, and further because corallicolids are typically present at low abundance. Metagenome-assembled genomes (MAGs) are commonly used to reconstruct genomes of uncultivated prokaryotes [79] and, more recently, eukaryotes [80]. However, MAG approaches are often less efficient for eukaryotes due to their large, repetitive genomes and tend to fail for low-abundant species in complex communities. Metagenomics-based transcriptomes (MGTs) therefore represent a valuable alternative, as they focus on the coding fraction of eukaryote genomes. This approach allows large-scale co-abundance clustering of transcripts across multiple samples to recover individual transcriptomes. To our knowledge, only one study has successfully applied this method, reconstructing 924 plankton transcriptomes that were largely inaccessible through MAGs approaches [81]. Here, by leveraging the extensive collection of *Pocillopora* metagenomes and metatranscriptomes, we reconstituted a corallicolid transcriptome. Although BUSCO analysis estimated the completeness of the corallicolid at only 33%, the here-generated resource constitutes a substantial addition to previous datasets generated using single-cell sequencing or density-gradient centrifugation [7]. Given the growing number of publicly available coral RNA-seq datasets, this study paves the way for more global reconstruction of the numerous eukaryotes associated to the large diversity of coral holobionts.

### Corallicolid diversity, abundance, and prevalence

To date, a large number of corallicolid 18S rRNA sequences have been reported (51 in the literature + 36 in this study) from only about 30 host species [5, 9, 16], suggesting the possibility that corallicolid diversity in anthozoans is underestimated. In *Pocillopora*, the presence of numerous rare and low-abundant 18SV9 variants as well as the co-occurrence of multiple ASVs within single colonies strongly suggest the existence of corallicolid intra-population diversity. Although we used the denoising tool DADA2 and strict filtering parameters, the presence of remaining sequencing errors that may inflate diversity estimates cannot be excluded. Nevertheless, the 6 dominant ASVs detected across many colonies likely represent distinct corallicolid populations.

Corallicolid apicomplexans are known to infect a broad range of hosts [5, 9]. Here, we show that single 18SV9 ASVs (ASV-1 and ASV-3) are present in all of the 6 *Pocillopora* species throughout the Pacific Ocean and are identical to sequences identified in anemones (*Metridium senile)* and corallimorpharians (Corynactis sp., *Rhodactis* sp.). This finding highlights the remarkable biogeographic and taxonomic breadth for specific corallicolid taxa. Long-distance dispersal via secondary hosts has recently been proposed as a potential mechanism explaining such distributions [82]. The substantial genetic divergence observed between available corallicolid transcriptomes (14% between Corallicolid ex. *Pocillopora* Corallicolid ex. *Parazoanthus*) suggests a long evolutionary history for this lineage. Additional transcriptomes and genomes will be required to uncover the genetic diversity and evolutionary trajectories of the corallicolid lineage.

We confirm that corallicolids are the second most abundant protist symbiont of corals and are nearly ubiquitous in *Pocillopora*. Although their average abundance remains low, exceptional cases were observed in which corallicolids exceeded 5% of ASV abundance, reaching levels comparable to Symbiodiniaceae ASVs, as previously reported [10]. Notably, corallicolids were also detected in colonies with low Symbiodiniaceae densities, indicating that the algal symbiont does not constrain the presence of the parasite, in agreement with earlier observations [15].

### Drivers of corallicolid abundance and gene expression

Although numerous abiotic parameters were measured *in situ* during the Tara expedition, no significant direct correlations were detected. Previous studies have shown that octocorals and hexacorals exposed to thermal stress harbor higher corallicolid abundances [15, 16]. In contrast, in our study, islands that previously experienced bleaching events did not exhibit higher corallicolid loads than other reefs. Moreover, *Pocillopora* colonies hosting *Durusdinium* rather than *Cladocopium*, did not show increased corallicolid abundance. Thus, corallicolid and Symbiodiniaceae diversity is not correlated.

Nevertheless, it is clear that corals from several reefs harbor substantially higher parasite loads than others. This spatial pattern may be linked to the coral species composition of each reef. For example, the corallicolid ASV-3 is significantly more abundant in *P. tuahiniensis*, a coral species sampled on Rapa Nui (I04) and Moorea (I07; Fig. 3A). These results suggests that corallicolid occurrence is more likely driven by its interactions with the coral host. The measurements of several coral physiological parameters on the coral colonies sequenced in this study revealed that ASV composition and gene expression patterns are partially explained by the rate of protein carbonylation and the oxidant scavenging capacities in the coral. The oxidative state of corals has been extensively studied because of its central role in the coral-Symbiodiniaceae symbiosis [83]. Although we did not detect a clear correlation between corallicolid abundance and the production of antioxidative compounds, we hypothesize that the ability of the coral host in maintaining a low level of ROS through efficient antioxidant defences is linked to the presence and activity of the corallicolids.

### Corallicolid gene content and host invasion strategy

Given that corallicolids belong to the phylum Apicomplexa, and are closely related to coccidians, it has long been assumed that they retain the canonical invasion mechanism of apicomplexans. However, until now, this assertion was based only on phylogenetic inference and their common ancestry with other apicomplexans, as no direct genomic evidence for apical organelles or invasion had been reported. Here, we provide an inventory of genes associated with the apicomplexan invasion machinery in corallicolids. All apicomplexans rely on micronemes and rhoptries to invade their hosts, and families of secretory genes are well characterized in model species such as *Toxoplasma* and *Plasmodium* [84, 85]. Genes encoding these core functions of apicomplexans were retrieved in the assembled corallicolid transcriptome and are highly expressed. Microscopy clearly indicates that corallicolids live in coral gastrodermal cells within a PV, however direct observation of their invasion, egress, or sexual stages are lacking [5]. We detected lineage-specific repertoires of PV-associated proteins, including a PV2 and GRA9 proteins. Despite limited sequence similarity, the putative corallicolid GRA9 identified here shares a conserved structural fold with apicomplexan GRA9 homologs, consistent with an evolutionary relationship and suggesting functional conservation. Several proteins of the glideosome were detected in the transcriptome. These proteins are required for motility, invasion and egress of *Toxoplasma gondii* and *Plasmodium falciparum.* Their expression in many samples reveal the active state of the corallicolids in *Pocillopora*.

### Corallicolid-coral crosstalk

The observation of corallicolids in visually healthy corals in this work and several previous studies has led to the suggestion that they are commensal members of the coral holobiont. In addition, our results highlight biological interactions indicative of an active crosstalk between the coral host and the parasite. Corallicolid gene expression patterns varied markedly among islands. In Rapa Nui island, we observed a higher corallicolid activity (replication, transcription, translation) consistent with a high relative abundance of this corallicolid in *Pocillopora* colonies. The low rate of coral carbonylation in this island and the low activity of corallicolid genes involved in the modulation of host immune defence, suggest that the host do not strongly react to the presence of the parasite. In contrast, in Hawaiian Islands, the upregulation of corallicolid genes involved in the modulation of host immune response is consistent with a stronger carbonylation and ROS scavenging capacities (TOSC) of the host. These two markers indicate a higher stress level for the coral potentially due to the activation of the immune system. These results suggest intense attack-defence dynamics in this island. Notably, these reefs also exhibited the highest relative abundance of corallicolids, reinforcing the hypothesis of a higher parasitic activity in these samples. While these patterns may reflect physiological differences between different *Pocillopora* species (*P.* tuahiniensis in Rapa Nui Island and *P. cf. effusa* or *P. meandrina* in Hawaiian Islands), we hypothesise that corallicolids around Hawaii elicit a stronger immune response from the host.

Overall, the strong expression of genes typically associated with parasitism together with differential regulation of pathways involved in host immune modulation and oxidative stress responses, support the view that corallicolids are active parasites that may influence coral immune system above an abundance threshold. Further studies focusing on the molecular and physiological interplay between corals and corallicolids will be essential to assess the potential role of these parasites in shaping coral health and resilience to environmental stressors.

## Acknowledgments

The Tara Pacific expedition would not have been possible without the participation and commitment of over 200 scientists, the Tara Ocean Foundation and its sailors, artists, staff and partners, the R/V Tara crew, and the Tara Pacific Expedition Participants [86]. We are keen to thank the commitment of the following institutions for their financial and scientific support that made this unique Tara Pacific Expedition possible: le Centre national de la recherche scientifique (CNRS), l’Université Paris Sciences & Lettres (PSL), le Centre Scientifique de Monaco (CSM), l’École Pratique des Hautes Études (EPHE), le Centre National de Séquençage - Genoscope, le Commissariat à l’énergie atomique et aux énergies alternatives (CEA), Inserm, l’Université Côte d’Azur, l’Agence Nationale de la Recherche (ANR), Agnès Troublé said agnès b., UNESCO-IOC, the Veolia Foundation, the Prince Albert II de Monaco Foundation, Région Bretagne, Billerudkorsnas, AmerisourceBergen Company, Lorient Agglomération, Oceans by Disney, L’Oréal, Biotherm, France Collectivités, Fonds Français pour l’Environnement Mondial (FFEM), Etienne Bourgois, and the Tara Ocean Foundation teams. Tara Pacific would not exist without the continuous support of the participating institutes. The authors also particularly thank Serge Planes, Denis Allemand, and the Tara Pacific consortium. This project was funded by France Génomique (ANR-10-INBS-09). We also acknowledge the Genoscope Sequencing Technical Team [87] and C. Scarpelli for support in high-performance computing. Computations were performed using the cobalt machine at the TGCC (CEA).

## Supplementary Figures

**Fig. S1:**
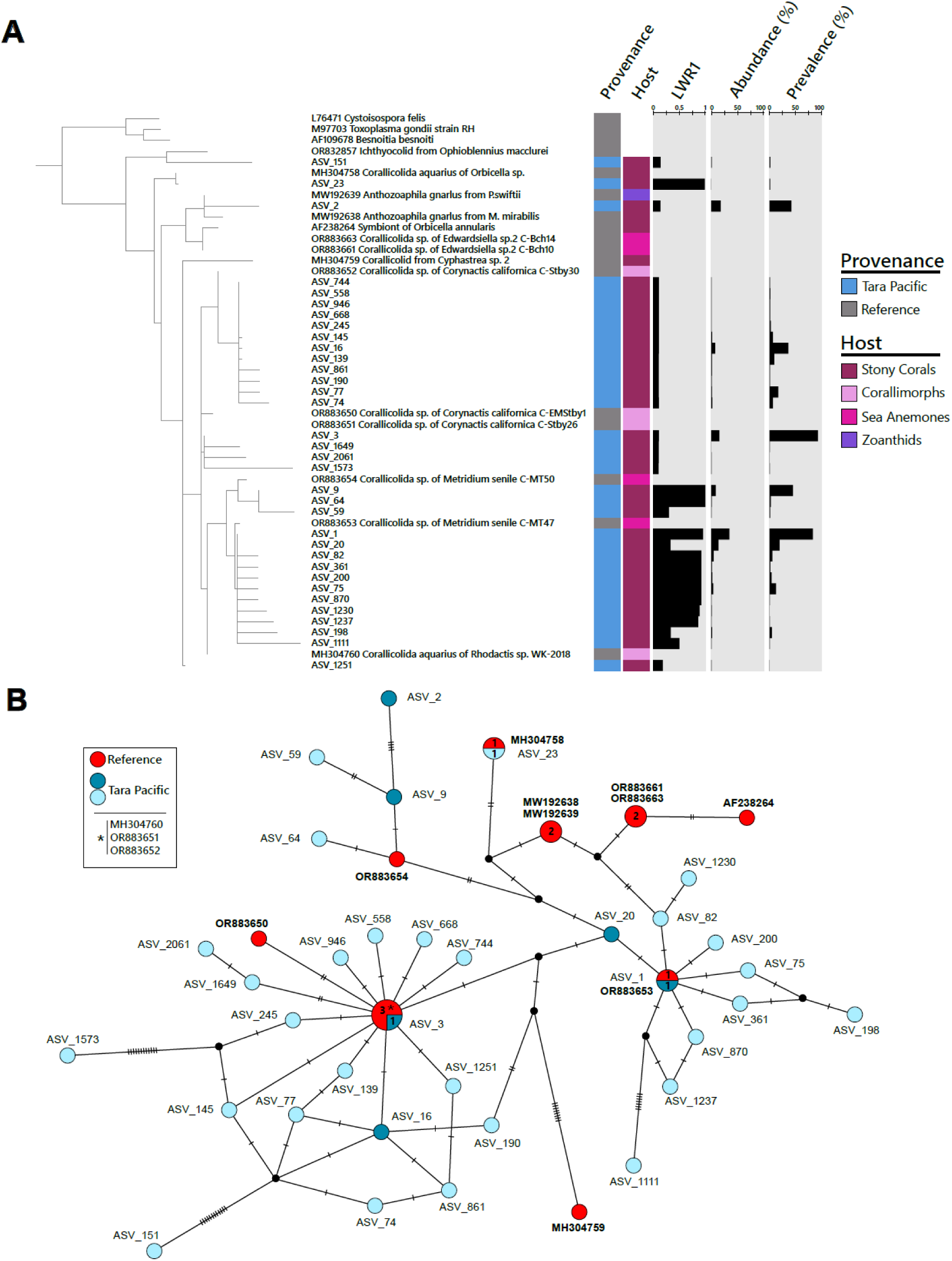
Phylogeny and TCS network of Corallicolid ASVs. A) Phylogenetic placement of corallicolid ASVs detected in Tara Pacific samples relative to reference corallicolid 18S rRNA sequences. A provenance track distinguishes environmental ASVs from reference sequences. Additional annotation tracks indicate the coral host type, ASV relative abundance, prevalence across samples, and the likelihood weight ratio of the primary placement (LWR1), reflecting placement confidence. B) Statistical parsimony (TCS) network of the 18S rRNA V9 region comparing Tara Pacific corallicolid ASVs with reference corallicolid sequences. Each node represents a unique V9 sequence variant; node size is proportional to the number of sequences sharing the same variant, with the number of sequences indicated inside the node when greater than one. Edges represent mutational steps, with hatch marks indicating the number of nucleotide differences. Black nodes correspond to inferred intermediate sequence variants (unsampled haplotypes) introduced by the TCS algorithm to connect observed variants parsimoniously. Node colors indicate sequence provenance, with reference sequences shown in red and Tara Pacific ASVs in blue. Among environmental ASVs, darker blue denotes the most abundant variants, while lighter blue corresponds to lower-abundance ASVs.

**Fig. S2:**
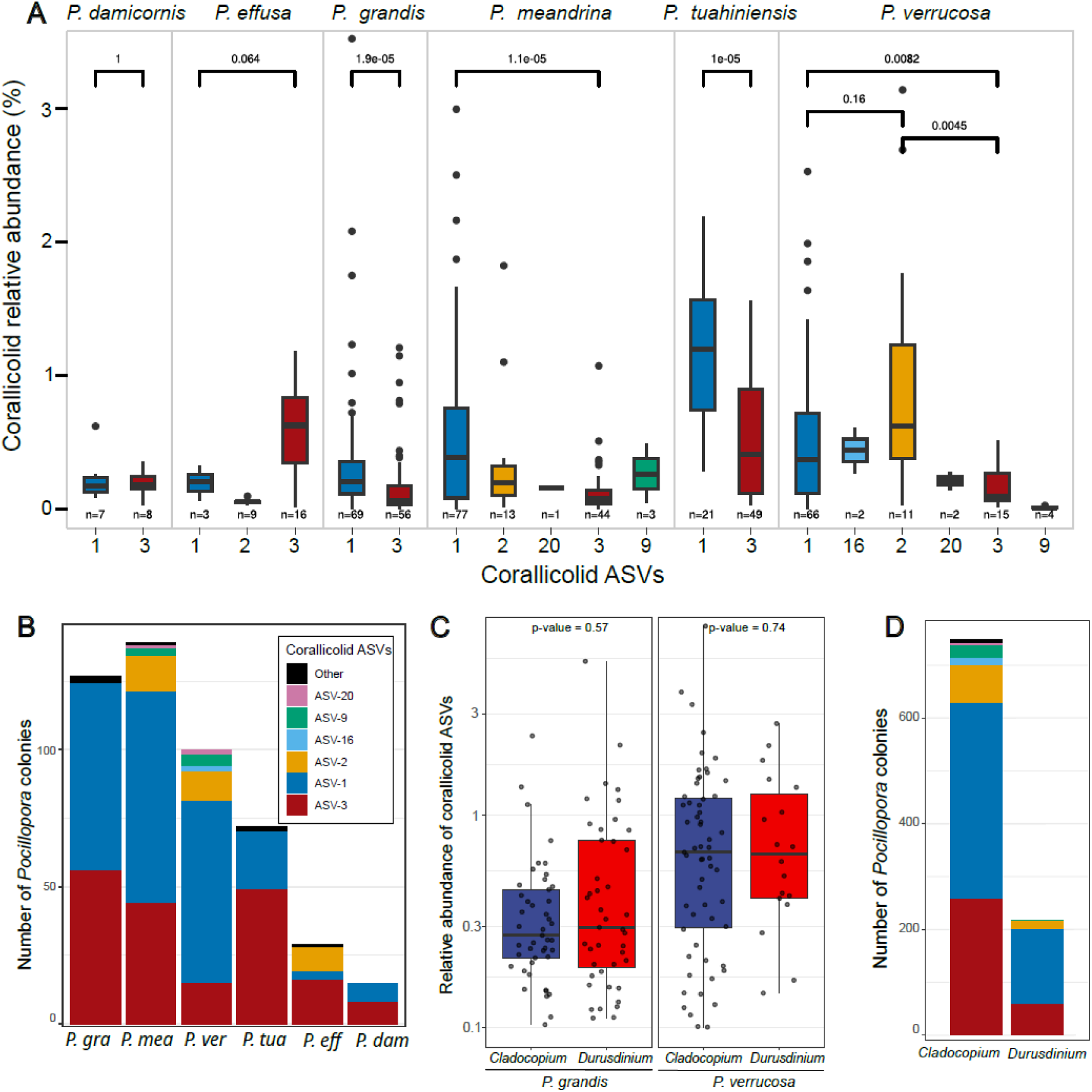
Prevalence and abundance of dominant corallicolid ASVs in *Pocillopora* colonies. A) Distribution of the relative abundance of the dominant corallicolid ASV in each *Pocillopora* species. p.values of wilcoxon statistical tests between the average abundance of ApiASV_1 and ASV-3 are indicated on each boxplot. B) Prevalence of most abundant Corallicolid ASV in each *Pocillopora* species. C) Distribution of corallicolid ASV relative abundance according to the presence of the thermosensitive *Cladocopium* or thermotolerant *Durusdinium* Symbiodiniaceae. Wilcoxon test p-values are indicated above each panel. D) Prevalence of most abundant corallicolid ASVs in *Pocillopora* colonies hosting each Symbiodiniaceae genus.

**Fig. S3:**
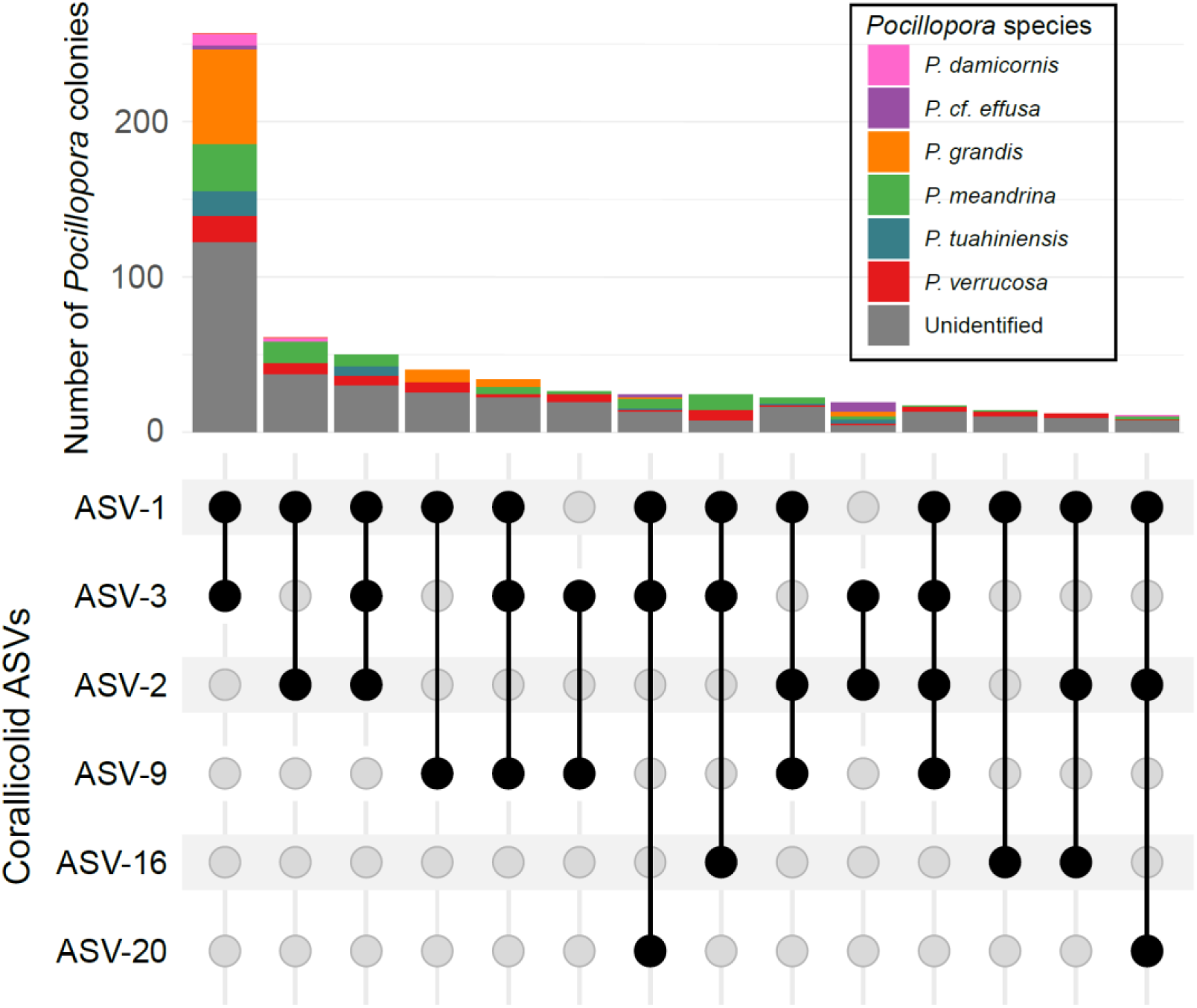
Corallicolid ASVs co-present in *Pocillopora* colonies. Upset plot representing the different ASV sequences detected within the same *Pocillopora* colonies. A corallicolid sequence was considered to be present if it represented more than 10% of all corallicolid ASVs.

**Fig. S4:**
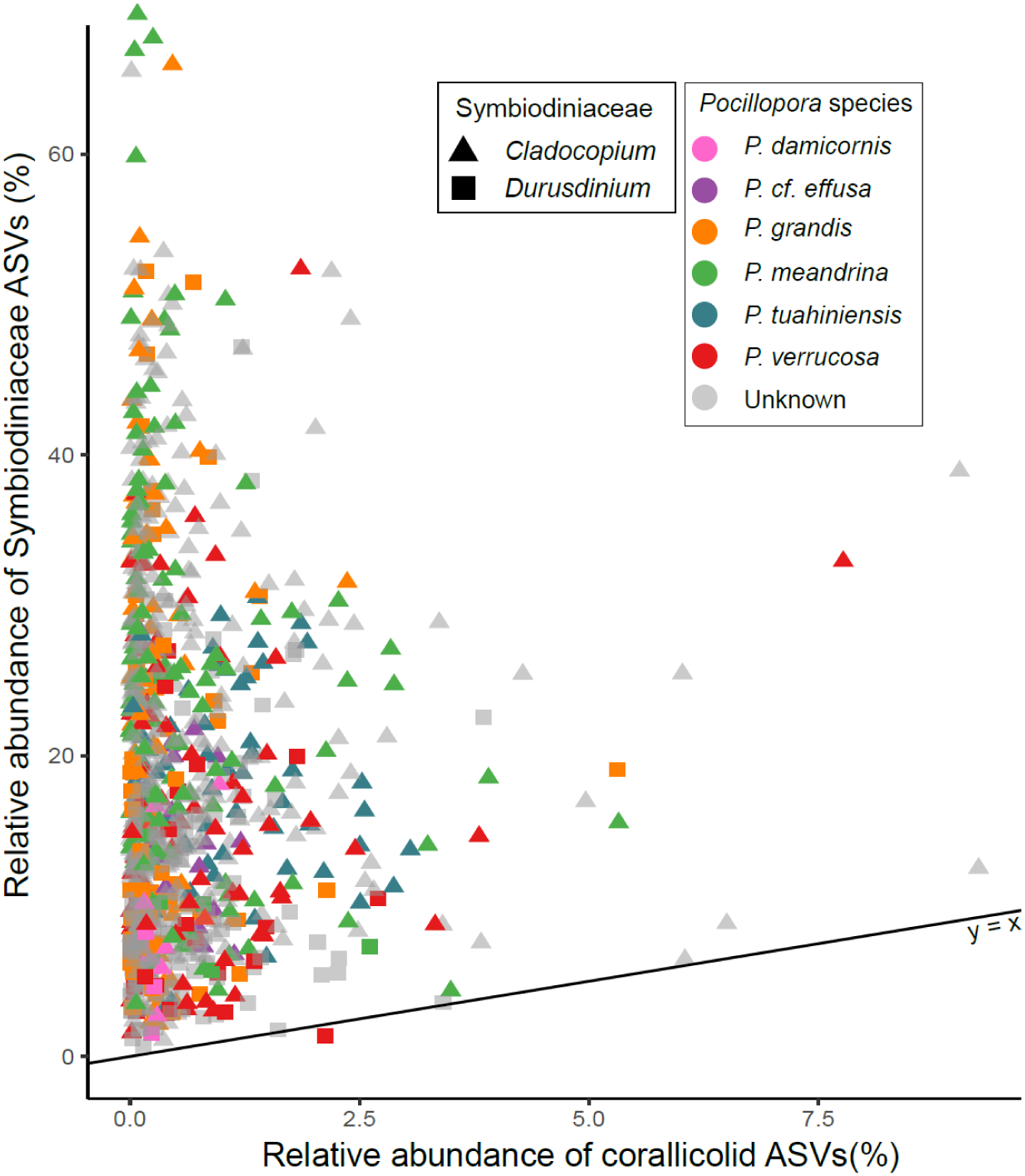
Relative abundance of corallicolids compared to Symbiodiniaceae. The proportion of corallicolid (x-axis) and Symbiodiniaceae (y-axis) 18S rRNA sequences relative to all eukaryote 18S rRNAs are represented for each *Pocillopora* sample. Dots are coloured by *Pocillopora* species and dot shapes indicate whether the symbiont belong to *Cladocopium* or *Durusdinium* genus. The black line represents the x=y function.

**Fig. S5:**
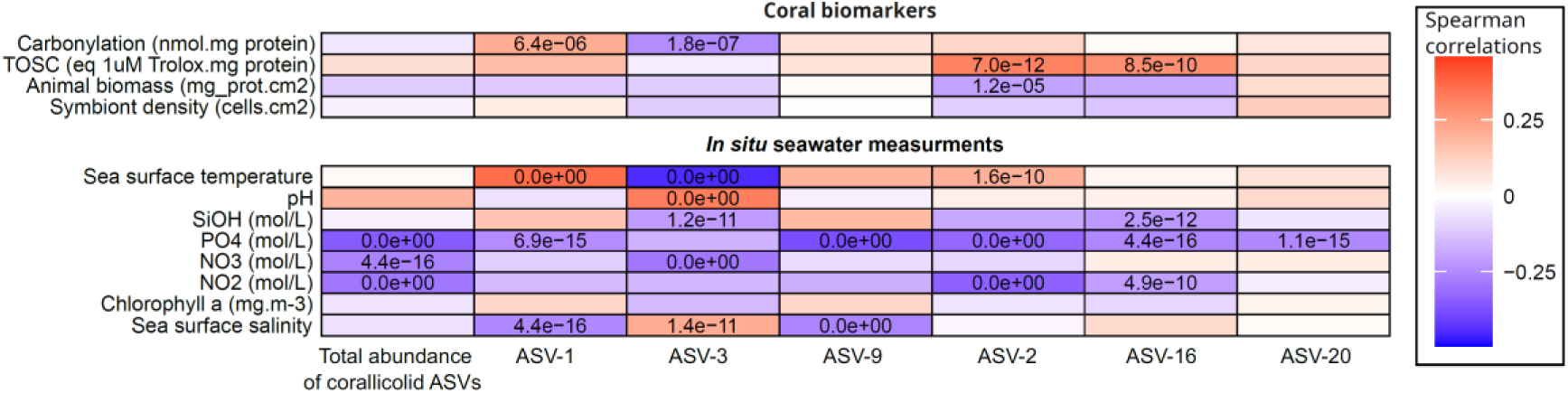
Correlations between corallicolid relative abundances and environmental parameters or coral biomarkers. Corallicolid relative abundances were calculated relatively to the total number of eukaryotes reads in each *Pocillopora* sample. Correlations were performed for all corallicolid ASVs or each ASV individually. Significant p-values < 1e-4 are indicated when Spearman correlations are > 0.2 or < -0.2.

**Fig. S6:**
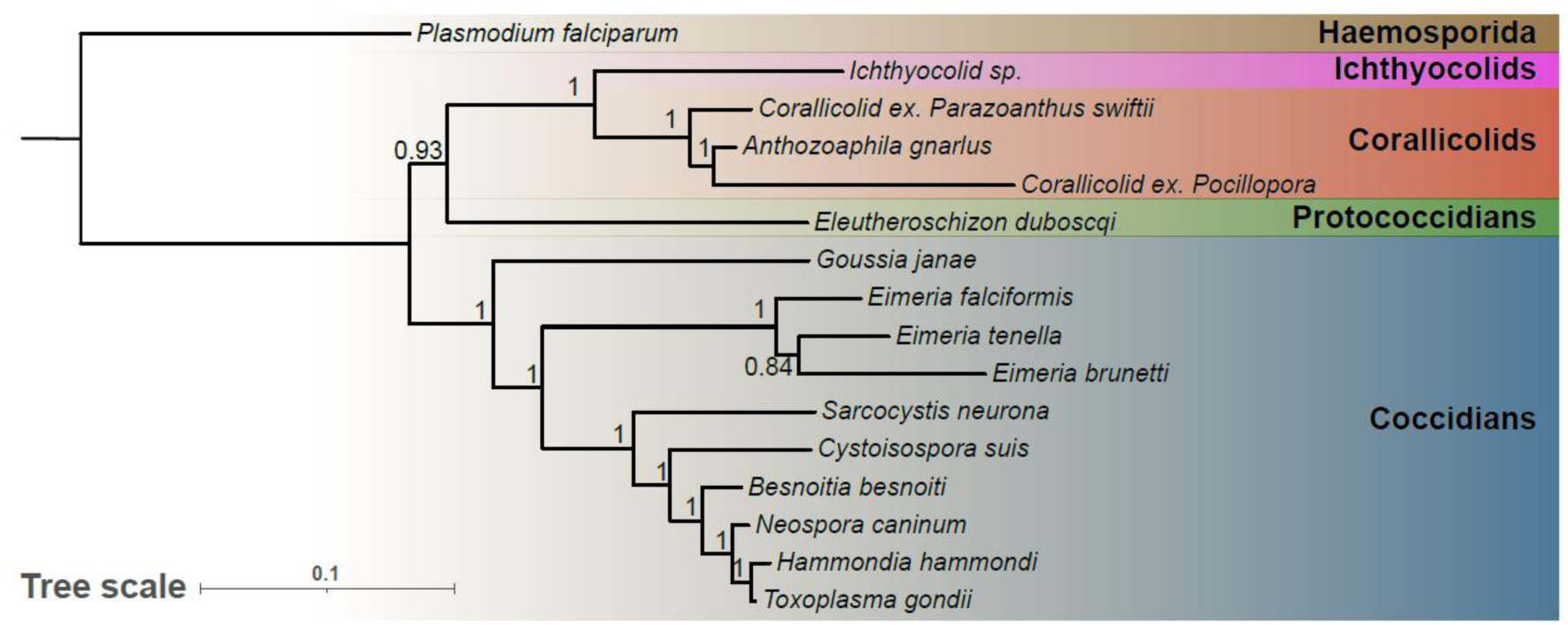
Phylogeny of Apicomplexa based on 102 core genes. The maximum-likelihood phylogeny was built from a multiple alignment of 102 proteins with 24,753 sites conserved across apicomplexans including the newly generated corallicolid transcriptome. GAMMA model of rate heterogeneity and LG substitution matrix were used with 100 bootstraps. Bootstrap values are indicated at each node.

**Fig. S7:**
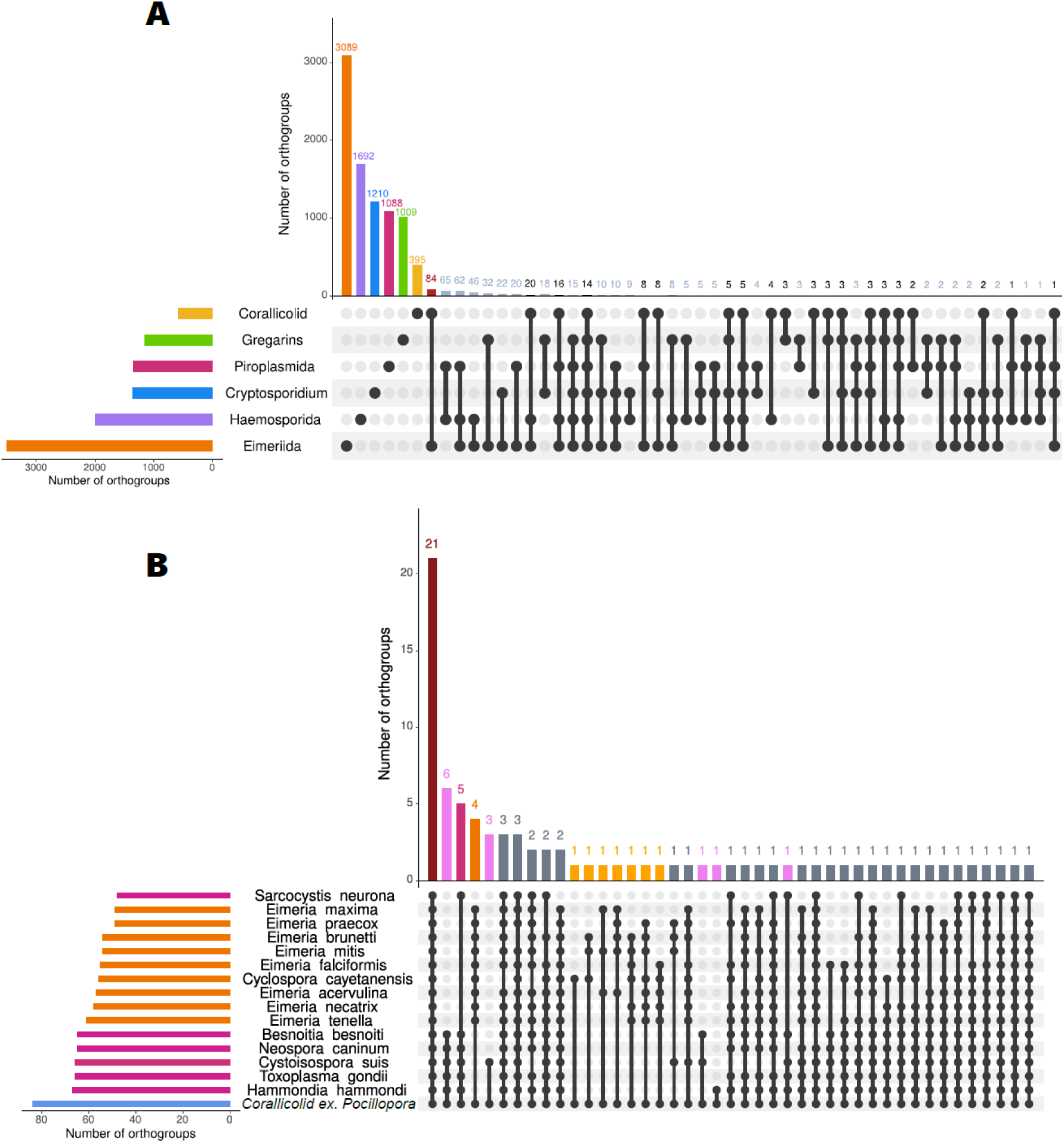
Orthology-based overview of corallicolid transcripts shared across parasitic Apicomplexa. Vertical bars indicate the number of orthogroups per intersection, while horizontal bars show the total number of orthogroups containing at least one protein from each clade. A) UpSet representation of orthogroups (OGs) restricted to parasitic apicomplexans, i.e. orthogroups in which at least one parasitic species was present and no proteins from non-parasitic chrompodellids (Chromera and Vitrella) were detected. Orthogroups containing at least one corallicolid protein are shown in black, whereas orthogroups lacking corallicolids are shown in grey. B) UpSet representation of the 84 orthogroups shared between corallicolids and Eimeriida, resolved at the species level. Eimeriida species are color-coded by family, with Eimeriidae shown in orange and Sarcocystidae in pink, corallicolid proteins are shown in blue. Intersections comprising all species within a given family are shown using the corresponding family color.

**Fig. S8:**
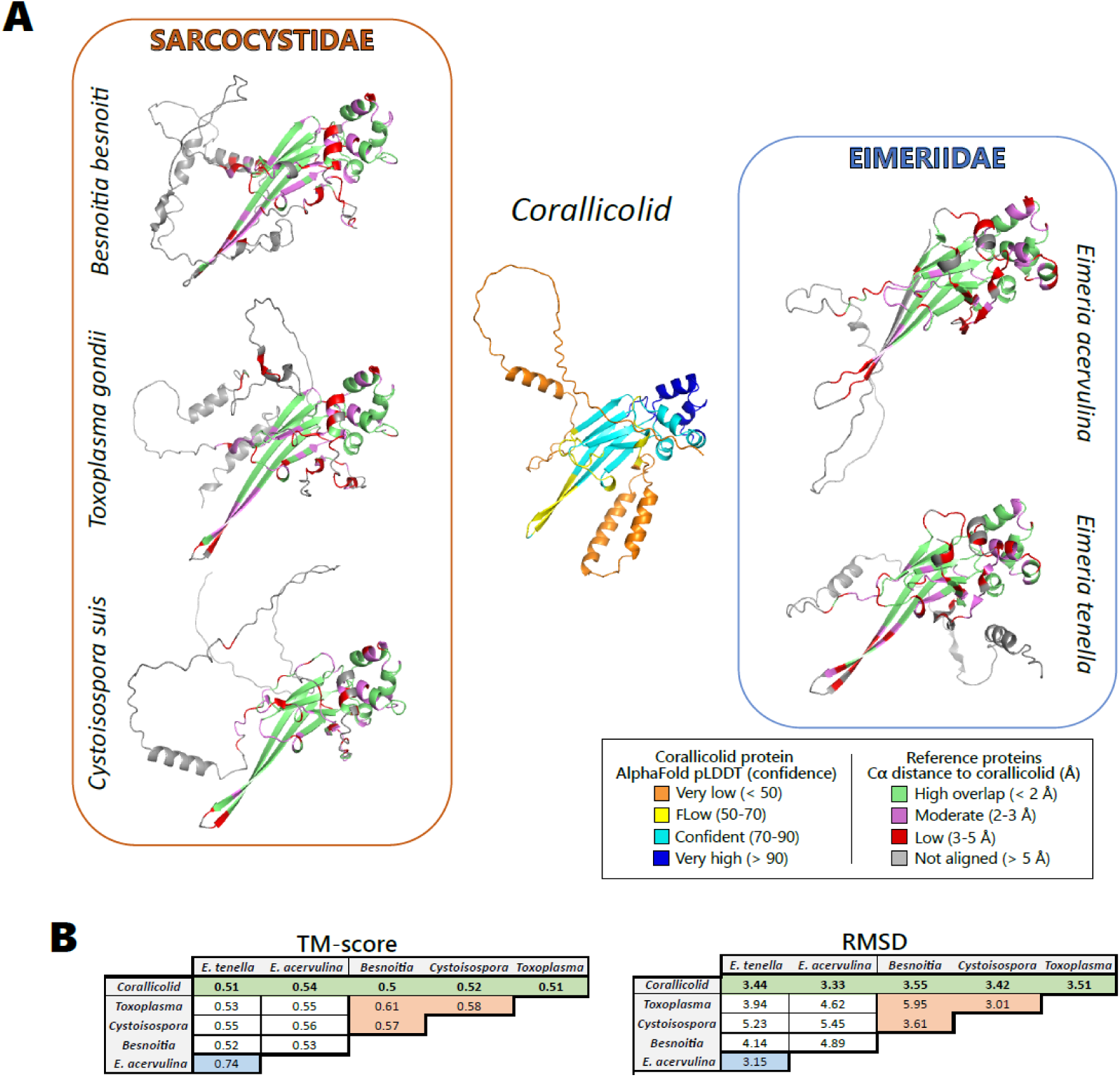
Structural conservation of GRA9 proteins. A) Structural comparison of selected GRA9 proteins from representative Sarcocystidae and Eimeriidae species against the corallicolid GRA9 AlphaFold model. The corallicolid structure is colored by pLDDT confidence, whereas reference structures are colored by Cα distance after superposition to indicate regions of high or low structural correspondence. B) Pairwise structural similarity is summarized using TM-scores and RMSD values (RMSD computed from structural alignments).

## References

1. Fisher R et al. Species Richness on Coral Reefs and the Pursuit of Convergent Global Estimates. Current Biology 2015;25:500–505. 10.1016/j.cub.2014.12.022

2. Hill TS, Hoogenboom MO. The indirect effects of ocean acidification on corals and coral communities. Coral Reefs 2022;41:1557–1583. 10.1007/s00338-022-02286-z

3. Hughes TP et al. Global warming transforms coral reef assemblages. Nature 2018;556:492–496. 10.1038/s41586-018-0041-2

4. Voolstra CR et al. Extending the natural adaptive capacity of coral holobionts. Nat Rev Earth Environ 2021;2:747–762. 10.1038/s43017-021-00214-3

5. Kwong WK et al. A widespread coral-infecting apicomplexan with chlorophyll biosynthesis genes. Nature 2019;568:103–107. 10.1038/s41586-019-1072-z

6. Bonacolta AM et al. A new and widespread group of fish apicomplexan parasites. Current Biology 2024;34:2748–2755.e3. 10.1016/j.cub.2024.04.084

7. Jacko-Reynolds VKL et al. Phylogenomics of coral-infecting corallicolids reveal multiple independent losses of chlorophyll biosynthesis in apicomplexan parasites. Current Biology 2025;35:1156–1163.e4. 10.1016/j.cub.2025.01.028

8. Jenkins BH, Waller RF. Parasite evolution: Coral-infecting apicomplexans disrupt our understanding of the pathways to parasitism. Current Biology 2025;35:R175–R177. 10.1016/j.cub.2024.12.050

9. Trznadel M et al. Coral-infecting parasites in cold marine ecosystems. Current Biology 2024;34:1810–1816.e4. 10.1016/j.cub.2024.03.026

10. Vohsen SA et al. Deep-sea corals provide new insight into the ecology, evolution, and the role of plastids in widespread apicomplexan symbionts of anthozoans. Microbiome 2020;8:34. 10.1186/s40168-020-00798-w

11. Janouškovec J et al. Global analysis of plastid diversity reveals apicomplexan-related lineages in coral reefs. Current Biology 2012;22:R518–R519. 10.1016/j.cub.2012.04.047

12. Kirk NL et al. Ubiquitous associations and a peak fall prevalence between apicomplexan symbionts and reef corals in Florida and the Bahamas. Coral Reefs 2013;32:847–858. 10.1007/s00338-013-1038-9

13. Mathur V et al. Global diversity and distribution of close relatives of apicomplexan parasites. Environmental Microbiology 2018;20:2824–2833. 10.1111/1462-2920.14134

14. Garcia GD et al. Metagenomic Analysis of Healthy and White Plague-Affected Mussismilia braziliensis Corals. Microb Ecol 2013;65:1076–1086. 10.1007/s00248-012-0161-4

15. Bonacolta AM et al. Differential apicomplexan presence predicts thermal stress mortality in the Mediterranean coral Paramuricea clavata. Environmental Microbiology 2024;26:e16548. 10.1111/1462-2920.16548

16. Peterson A et al. Apicomplexan and non-metazoan microeukaryotes in the thermosensitive reef-building coral Acropora hyacinthus shift in abundance throughout an extreme coral bleaching event. Front Mar Sci 2025;12. 10.3389/fmars.2025.1626071

17. Planes S et al. The Tara Pacific expedition—A pan-ecosystemic approach of the “-omics” complexity of coral reef holobionts across the Pacific Ocean. PLOS Biology . 2019. 2019., 17: e3000483

18. Lombard F et al. Open science resources from the Tara Pacific expedition across coral reef and surface ocean ecosystems. Scientific Data . 2023. 2023., 10: 324

19. Belser C et al. Integrative omics framework for characterization of coral reef ecosystems from the Tara Pacific expedition. 2022. 10.48550/arXiv.2207.02475

20. Henry N et al. rDNA 18S V9 ASVs (DADA2) from the Tara Pacific Expedition. 2024. Zenodo, 2024.

21. Hume BCC et al. Tara Pacific CDIV ITS2 Symbiodiniaceae data release. 2022. Zenodo, 2022.

22. Deshuraud R et al. From genome wide SNPs to genomic islands of differentiation: the quest for species diagnostic markers in two scleractinian corals, Pocillopora and Porites. 2022;2022.10.21.513203. 10.1101/2022.10.21.513203

23. Noel B et al. Pervasive tandem duplications and convergent evolution shape coral genomes. Genome Biol 2023;24:123. 10.1186/s13059-023-02960-7

24. Voolstra CR et al. Disparate genetic divergence patterns in three corals across a pan-Pacific environmental gradient highlight species-specific adaptation. npj biodivers 2023;2:1–16. 10.1038/s44185-023-00020-8

25. Porro B et al. Different environmental response strategies in sympatric corals from Pacific Islands. Commun Earth Environ 2023;4:311. 10.1038/s43247-023-00946-8

26. Zamoum T, Furla P. Symbiodinium isolation by NaOH treatment. J Exp Biol 2012;215:3875–3880. 10.1242/jeb.074955

27. Buss H et al. Protein Carbonyl Measurement by a Sensitive ELISA Method. Free Radical Biology and Medicine 1997;23:361–366. 10.1016/S0891-5849(97)00104-4

28. Naguib YMA. A Fluorometric Method for Measurement of Oxygen Radical-Scavenging Activity of Water-Soluble Antioxidants. Analytical Biochemistry 2000;284:93–98. 10.1006/abio.2000.4691

29. Guillou L et al. The Protist Ribosomal Reference database (PR2): a catalog of unicellular eukaryote Small Sub-Unit rRNA sequences with curated taxonomy. Nucleic Acids Res 2013;41:D597–D604. 10.1093/nar/gks1160

30. Murali A, Bhargava A, Wright ES. IDTAXA: a novel approach for accurate taxonomic classification of microbiome sequences. Microbiome 2018;6:140. 10.1186/s40168-018-0521-5

31. Rognes T et al. VSEARCH: a versatile open source tool for metagenomics. PeerJ 2016;4:e2584. 10.7717/peerj.2584

32. Clement M, Posada D, Crandall KA. TCS: a computer program to estimate gene genealogies. Molecular Ecology 2000;9:1657–1659. 10.1046/j.1365-294x.2000.01020.x

33. Leigh JW, Bryant D. popart: full-feature software for haplotype network construction. Methods in Ecology and Evolution 2015;6:1110–1116. 10.1111/2041-210X.12410

34. Chen X-H et al. Quantitative Proteomics Reveals Common and Specific Responses of a Marine Diatom Thalassiosira pseudonana to Different Macronutrient Deficiencies. Front Microbiol 2018;9:2761. 10.3389/fmicb.2018.02761

35. Dougan KE et al. Whole-genome duplication in an algal symbiont bolsters coral heat tolerance. Science Advances 2024;10:eadn2218. 10.1126/sciadv.adn2218

36. Li H et al. The Sequence Alignment/Map format and SAMtools. Bioinformatics 2009;25:2078–2079. 10.1093/bioinformatics/btp352

37. Fu L et al. CD-HIT: accelerated for clustering the next-generation sequencing data. Bioinformatics 2012;28:3150–3152. 10.1093/bioinformatics/bts565

38. Haas BJ et al. TransDecoder. 2026. TransDecoder, 2026.

39. Buchfink B, Reuter K, Drost H-G. Sensitive protein alignments at tree-of-life scale using DIAMOND. Nat Methods 2021;18:366–368. 10.1038/s41592-021-01101-x

40. Groussman RD et al. MarFERReT, an open-source, version-controlled reference library of marine microbial eukaryote functional genes. Sci Data 2023;10:926. 10.1038/s41597-023-02842-4

41. Camacho C, et al. BLAST+: architecture and applications. BMC Bioinformatics 2009;10:421. 10.1186/1471-2105-10-421

42. Alneberg J et al. Binning metagenomic contigs by coverage and composition. Nat Methods 2014;11:1144–1146. 10.1038/nmeth.3103

43. Simão FA et al. BUSCO: assessing genome assembly and annotation completeness with single-copy orthologs. Bioinformatics 2015;31:3210–3212. 10.1093/bioinformatics/btv351

44. Aramaki T et al. KofamKOALA: KEGG Ortholog assignment based on profile HMM and adaptive score threshold. Bioinformatics 2020;36:2251–2252. 10.1093/bioinformatics/btz859

45. Quevillon E et al. InterProScan: protein domains identifier. Nucleic Acids Res 2005;33:W116–120. 10.1093/nar/gki442

46. Katoh K, Standley DM. MAFFT Multiple Sequence Alignment Software Version 7: Improvements in Performance and Usability. Mol Biol Evol 2013;30:772–780. 10.1093/molbev/mst010

47. Kozlov AM et al. RAxML-NG: a fast, scalable and user-friendly tool for maximum likelihood phylogenetic inference. Bioinformatics 2019;35:4453–4455. 10.1093/bioinformatics/btz305

48. Berger SA, Stamatakis A. PaPaRa 2.0: a vectorized algorithm for probabilistic phylogeny-aware alignment extension. Heidelberg Institute for Theoretical Studies 2012;12.

49. Barbera P et al. EPA-ng: Massively Parallel Evolutionary Placement of Genetic Sequences. Syst Biol 2019;68:365–369. 10.1093/sysbio/syy054

50. Czech L, Barbera P, Stamatakis A. Genesis and Gappa: processing, analyzing and visualizing phylogenetic (placement) data. Bioinformatics 2020;36:3263–3265. 10.1093/bioinformatics/btaa070

51. Eren AM et al. Anvi’o: an advanced analysis and visualization platform for ‘omics data. PeerJ 2015;3:e1319. 10.7717/peerj.1319

52. Capella-Gutiérrez S, Silla-Martínez JM, Gabaldón T. trimAl: a tool for automated alignment trimming in large-scale phylogenetic analyses. Bioinformatics 2009;25:1972–1973. 10.1093/bioinformatics/btp348

53. Lemoine F et al. Renewing Felsenstein’s phylogenetic bootstrap in the era of big data. Nature 2018;556:452–456. 10.1038/s41586-018-0043-0

54. Letunic I, Bork P. Interactive Tree of Life (iTOL) v6: recent updates to the phylogenetic tree display and annotation tool. Nucleic Acids Res 2024;52:W78–W82. 10.1093/nar/gkae268

55. Emms DM, Kelly S. OrthoFinder: phylogenetic orthology inference for comparative genomics. Genome Biol 2019;20:238. 10.1186/s13059-019-1832-y

56. Amos B et al. VEuPathDB: the eukaryotic pathogen, vector and host bioinformatics resource center. Nucleic Acids Res 2021;50:D898–D911. 10.1093/nar/gkab929

57. Jones P et al. InterProScan 5: genome-scale protein function classification. Bioinformatics 2014;30:1236–1240. 10.1093/bioinformatics/btu031

58. Bailey TL, et al. The MEME Suite. Nucleic Acids Res 2015;43:W39–W49. 10.1093/nar/gkv416

59. Lemoine F, Gascuel O. Gotree/Goalign: toolkit and Go API to facilitate the development of phylogenetic workflows. NAR Genom Bioinform 2021;3:lqab075. 10.1093/nargab/lqab075

60. Nguyen L-T et al. IQ-TREE: A Fast and Effective Stochastic Algorithm for Estimating Maximum-Likelihood Phylogenies. Mol Biol Evol 2015;32:268–274. 10.1093/molbev/msu300

61. Abramson J et al. Accurate structure prediction of biomolecular interactions with AlphaFold 3. Nature 2024;630:493–500. 10.1038/s41586-024-07487-w

62. Schrödinger, LLC. The PyMOL Molecular Graphics System, Version 1.8. 2015. 2015.

63. Zhang Y, Skolnick J. TM-align: a protein structure alignment algorithm based on the TM-score. Nucleic Acids Res 2005;33:2302–2309. 10.1093/nar/gki524

64. Mayer K-I et al. Tara Pacific CDIV ITS2 Symbiodiniaceae data release V2. 2025. Zenodo, 2025.

65. Briquet S et al. High-Mobility-Group Box Nuclear Factors of Plasmodium falciparum. Eukaryot Cell 2006;5:672–682. 10.1128/EC.5.4.672-682.2006

66. Komaki-Yasuda K, Kano S. The RNA-binding KH-domain in the unique transcription factor of the malaria parasite is responsible for its transcriptional regulatory activity. PLoS One 2023;18:e0296165. 10.1371/journal.pone.0296165

67. Painter HJ, Campbell TL, Llinás M. The Apicomplexan AP2 family: Integral factors regulating *Plasmodium* development. Molecular and Biochemical Parasitology 2011;176:1–7. 10.1016/j.molbiopara.2010.11.014

68. Saharan K et al. Structure-function studies of a nucleoplasmin isoform from Plasmodium falciparum. J Biol Chem 2025;301:108379. 10.1016/j.jbc.2025.108379

69. Kono M et al. The apicomplexan inner membrane complex. Front Biosci (Landmark Ed) 2013;18:982–992. 10.2741/4157

70. Dobson SE et al. The crystal structures of macrophage migration inhibitory factor from Plasmodium falciparum and Plasmodium berghei. Protein Science 2009;18:2578–2591. 10.1002/pro.263

71. Sommerville C et al. Biochemical and Immunological Characterization of *Toxoplasma gondii* Macrophage Migration Inhibitory Factor*. Journal of Biological Chemistry 2013;288:12733–12741. 10.1074/jbc.M112.419911

72. Bao J et al. Serpin functions in host-pathogen interactions. PeerJ 2018;6:e4557. 10.7717/peerj.4557

73. Mansouri R et al. Cyclophilins as key players in protozoan parasite infections. Parasites Vectors 2025;18:457. 10.1186/s13071-025-07098-y

74. Tampaki Z et al. Ectopic Expression of a Neospora caninum Kazal Type Inhibitor Triggers Developmental Defects in Toxoplasma and Plasmodium. PLOS ONE 2015;10:e0121379. 10.1371/journal.pone.0121379

75. Lian L et al. Inhibition of Cell Apoptosis by Apicomplexan Protozoa–Host Interaction in the Early Stage of Infection. Animals 2023;13. 10.3390/ani13243817

76. Keeling PJ, Eglit Y. Openly available illustrations as tools to describe eukaryotic microbial diversity. PLOS Biology 2023;21:e3002395. 10.1371/journal.pbio.3002395

77. Adjogble KDZ et al. GRA9, a new *Toxoplasma gondii* dense granule protein associated with the intravacuolar network of tubular membranes. International Journal for Parasitology 2004;34:1255–1264. 10.1016/j.ijpara.2004.07.011

78. Toller W, Rowan R, Knowlton N. Genetic evidence for a protozoan (phylum Apicomplexa) associated with corals of the Montastraea annularis species complex. Coral Reefs 2002;21:143–146. 10.1007/s00338-002-0220-2

79. Wu D et al. A metagenomic perspective on the microbial prokaryotic genome census. Science Advances 2025;11:eadq2166. 10.1126/sciadv.adq2166

80. Delmont TO et al. Functional repertoire convergence of distantly related eukaryotic plankton lineages abundant in the sunlit ocean. Cell Genomics 2022;2:100123. 10.1016/j.xgen.2022.100123

81. Vorobev A et al. Transcriptome reconstruction and functional analysis of eukaryotic marine plankton communities via high-throughput metagenomics and metatranscriptomics. Genome Res 2020;30:647–659. 10.1101/gr.253070.119

82. Bonacolta AM et al. Fireworms are a reservoir and potential vector for coral-infecting apicomplexans. ISME J 2025;19:wraf078. 10.1093/ismejo/wraf078

83. Bhattacharya D et al. The Host Coral Bleaching Response Viewed Through the Lens of Multi-Omics. BioEssays 2026;48:e70110. 10.1002/bies.70110

84. Dubois DJ, Soldati-Favre D. Biogenesis and secretion of micronemes in Toxoplasma gondii. Cellular Microbiology 2019;21:e13018. 10.1111/cmi.13018

85. Mageswaran SK et al. In situ ultrastructures of two evolutionarily distant apicomplexan rhoptry secretion systems. Nat Commun 2021;12:4983. 10.1038/s41467-021-25309-9

86. Tara Pacific Expedition Participants. Tara Pacific Expedition Participants. 2020. 10.5281/zenodo.3777760

87. Oliveira PH. Genoscope Sequencing Technical Team (GSTT) author list. 2025. 10.5281/zenodo.14611490

